# Canonical Hidden Markov Model Networks for Studying M/EEG

**DOI:** 10.1101/2025.10.21.683692

**Authors:** Chetan Gohil, Rukuang Huang, Cameron Higgins, Mats W.J. van Es, Andrew J. Quinn, Diego Vidaurre, Mark W. Woolrich

## Abstract

Dynamic brain networks identified in magneto/encephalography (M/EEG) recordings provide new insights into human brain activity. One established method uses Hidden Markov Models (HMMs) and has been shown to infer reproducible, fast-switching brain networks in a variety of cognitive and disease conditions. Often these studies are done on small bespoke (*boutique*) datasets (*N* < 100) and analysed in isolation of other M/EEG datasets. Instead of training a new model for each boutique study, which is computationally expensive, we propose the use of a *canonical HMM*. This provides a common reference through which different studies can be described using the same set of networks. We provide HMMs for a range of model orders (4-16 states) in parcellated source space and sensor space. These HMMs were trained on 1849 MEG recordings (*N* = 621, 18-88 years old, 194 hours), capturing population variability in both rest and task data. We illustrate applications of this canonical HMM approach in parcellated source space using boutique MEG and EEG datasets. Applying the canonical HMM in parcel space requires the boutique dataset to be preprocessed and source reconstructed in the same way as the canonical HMM training data. Applying the canonical HMM in the sensor space requires the same sensor layout and preprocessing as the canonical HMM training data. The canonical HMMs have been made publicly available as an open-access resource, providing sets of canonical brain networks that can be used to compare individuals within and across a range of datasets.

**Highlights:**

- Instead of training a new HMM for every study, which is computationally expensive and can lead to less reliable results, we propose a *canonical HMM* instead.
- The canonical HMM provides a common reference through which different studies can be described using the same set of networks.
- The canonical HMM was learnt from 1849 MEG recordings (194 hours of data; resting state and task) in participants aged 18–88 years.
- We demonstrate applications of the canonical HMM on three parcellated source space boutique studies: an Alzheimer’s resting-state MEG dataset, a working memory task MEG dataset, and a resting-state EEG dataset.
- We transfer the parcel-level canonical HMM to sensor space, avoiding the need for source reconstruction.
- We release the canonical HMM (for varying model orders) as an open-access resource for M/EEG research.

## 1 Introduction

Cortical activity, as measured by electrophysiological recordings, can be described using transient, re-occurring large-scale networks of phase-locking (coherent) activity (Vidaurre et al., 2018b). These networks, which exhibit fast dynamics (∼ 100 ms) and correspond to transient oscillatory activity across different regions (*spatio-spectral patterns*), are typically estimated using a Hidden Markov Model (HMM) applied to source-reconstructed magneto/encephalography (M/EEG) data (Rabiner and Juang, 2003; Bishop and Nasrabadi, 2006; Cho et al., 2024). The networks and their dynamics are learnt directly from the data, usually without the user specifying the spatio-spectral patterns. See (Gohil et al., 2024a) for a description of the HMM approach applied to M/EEG data and Section 2 for further details.

The transient network perspective provided by the HMM has been used to gain an understanding of cognition (Guichet et al., 2025; Rossi et al., 2023; Fauchon et al., 2022; Higgins et al., 2021) and disease (Kohl et al., 2025; Seedat et al., 2023; Sitnikova et al., 2018). To gain specific insights, HMM analysis is often performed on bespoke datasets; for example, by looking at the network-level responses to tasks (Gohil et al., 2024b), or by contrasting network characteristics between healthy and disease groups (Kohl et al., 2025; Seedat et al., 2023; Sitnikova et al., 2018). In this work, we will refer to these as *boutique* datasets. However, the relatively small size of these datasets (typically less than ∼100 participants) often reduces the detail and reliability of the estimated large-scale networks and their dynamics (discussed further in Section 4.7).

The HMM has previously been applied successfully to several independent boutique M/EEG datasets (Vidaurre et al., 2018b; Cho et al., 2024; Gohil et al., 2024a; Guichet et al., 2025; Rossi et al., 2023; Fauchon et al., 2022; Higgins et al., 2021; Kohl et al., 2025; Seedat et al., 2023; Sitnikova et al., 2018; Gohil et al., 2024b; van Es et al., 2023; Gohil et al., 2022; Quinn et al., 2018; Vidaurre et al., 2018a, 2016; Coquelet et al., 2022; Zhang et al., 2021). Broadly, the HMM applied to M/EEG data finds a consistent set of networks across a variety of tasks and at rest (Gohil et al., 2024a). A similar consistency has also been observed in the functional networks found using functional magnetic resonance imaging (fMRI) (Smith et al., 2009). There is also generally good correspondence between networks found in MEG and EEG (Cho et al., 2024) and between healthy and diseased populations (Kohl et al., 2025; Sitnikova et al., 2018).

Given the commonality, we propose adopting a *canonical* reference model, rather than training a new HMM for each dataset in isolation. In this work, we present a canonical HMM pre-trained on a large MEG dataset, which provides a set of networks that describe re-occurring spatio-spectral patterns of activity in electrophysiological (MEG and EEG) data. In a similar spirit to the *Yeo atlas* (or *parcellation*) of large-scale functional networks derived from resting-state fMRI (Yeo et al., 2011), we propose that M/EEG research would benefit from a standardised set of dynamic electrophysiological networks. The Yeo atlas has facilitated reproducibility and comparability across diverse fMRI datasets by offering a common parcellation of the cortex based on functional connectivity. Analogously, the canonical HMM provides a dynamic network reference for M/EEG.

The canonical HMM has some key benefits: it provides a basis set of networks that can be used to compare different studies, including individuals or groups, in a common reference space; it improves the stability of network inference; and lowers the computational resources required to perform an HMM analysis on new data.

In this work, we have developed a canonical HMM framework that includes a set of pre-trained models with a different number of states (4 to 16) in both parcellated source space and in MEG sensor space. The main constraint in applying these models is that the new datasets have the same spatial layout/parcellation and are subject to similar preprocessing/source reconstruction to that used in the canonical HMM (described in Section 2 and discussed further in Section 4).

To pre-train the canonical HMM, we have used the Cam-CAN dataset (see Section 2.1.1), which contains recordings from a large healthy population (*N* = 621, 1849 rest and task sessions). This is an ideal dataset due to its size (one of the largest currently available) and its wide age range of participants (18-88 years old). The canonical HMMs, along with example code for applying them to new datasets, have been made available as a public resource, see Section 6 for further details.

## 2 Materials and Methods

### 2.1 Datasets

#### 2.1.1 Training MEG Dataset: Cam-CAN (Resting State and Task, 1849 Sessions)

To train the canonical HMM, we use a group of 621 healthy participants aged between 18 and 88 years from the Cam-CAN (Cambridge Centre for Ageing and Neuroscience) dataset (Shafto et al., 2014; Taylor et al., 2017). Each participant has multiple Elekta MEG recordings: an eyes closed resting-state session; a sensorimotor task session; and a passive visual/auditory task session. A total of 1849 sessions were studied in this work. The demographics of the participants studied are shown in Figure S1A. Further information regarding the dataset, protocols, tasks and the MEG/MRI acquisition is provided in (Shafto et al., 2014; Taylor et al., 2017).

#### 2.1.2 Example MEG Boutique Dataset: BioFIND (Resting State, 82 Sessions)

In this work, we studied a subset of the BioFIND dataset (Hughes et al., 2019; Vaghari et al., 2022), which aims to identify biological markers for the early detection of Alzheimer’s disease. We analysed data from the Cambridge site and excluded those with missing electrooculography recordings or sMRIs. This resulted in 41 MCI participants and 41 healthy controls, each with a 4-13 minute eyes closed resting-state Elekta MEG recording. The demographics of the participants studied are shown in Figure S1B. Further information regarding this dataset, the participants, protocols and the MEG/MRI acquisition can be found in (Hughes et al., 2019; Vaghari et al., 2022).

#### 2.1.3 Example MEG Boutique Dataset: MEGUK (*N*-back Task, 74 Sessions)

In this work, we studied a subset of the MEGUK partnership dataset (MEGUK Partnership, 2025), which is large collaborative effort to share MEG data from multiple sites. We analysed data from the Cardiff site excluding participants who did not have head shape digitisation points. We studied 74 CTF MEG recordings from healthy participants performing a working memory task: a visual *N*-back task (Owen et al., 2005) with 0-back, 1-back, and 2-back conditions. The demographics of the participants studied are shown in Figure S1C. Further information regarding this dataset can be found in (MEGUK Partnership, 2025).

#### 2.1.4 Example EEG Boutique Dataset: LEMON (Resting State, 108 Sessions)

In this work, we studied a subset of the LEMON (Leipzig Study for Mind-Body-Emotion Interactions) dataset (Babayan et al., 2019) excluding any participant in the full dataset that did not have a structural MRI or Polhemus head shape points. This resulted in 108 healthy participants. Medium density (61 channel) EEG data were recorded from each participant in a resting-state session where they alternated between periods of eyes closed and eyes open every 30 seconds. The demographics of the participants studied are shown in Figure S1D. Further information regarding this dataset, protocols and the MEG/MRI acquisition can be found in (Babayan et al., 2019).

### 2.2 M/EEG Preprocessing, Source Reconstruction and Parcellation

Each dataset was processed using the osl-ephys toolbox (van Es et al., 2025; Quinn et al., 2022). For each session, we applied the following steps^1^:

1. **Filtering and downsampling.** First, we bandpass filter the data (forward-backward fifth-order IIR Butterworth) between 0.5 and 125 Hz, then apply notch filters at 50 Hz and 100 Hz to remove mains artefacts. For Cam-CAN an additional spike artefact at 88 Hz was also removed using a notch filter. Following this, the data were downsampled to 250 Hz.
2. **Automated bad segment/channel detection.** Bad segments and channels can have a detrimental effect on independent component analysis (ICA) denoising (next step). Therefore, we annotated significantly high variance (*p*-value < 0.1) using an automated generalised-extreme studentised deviate (G-ESD) algorithm (Rosner, 1983). Bad segment and channel detection was applied separately to the different sensor types (magnetometers, gradiometers, EEG).
3. **ICA artefact removal.** After excluding the bad segments and channels, we applied FastICA (Hyvarinen, 1999) to the sensor-level MEG data, decomposing the signal into 64 independent components. For the LEMON dataset, we decomposed the sensor-level EEG into 30 independent components. Based on the correlation of the ICA component time courses with the EOG/ECG electrodes (0.9 threshold), independent components were marked as ocular/cardiac noise and removed from the MEG data.
4. **Bad channel interpolation.** Bad channels in the ICA-cleaned M/EEG data were replaced using a spherical spline interpolation (Perrin et al., 1989). This was the final step in preprocessing the sensor-level MEG data.
5. **EEG re-referencing.** The EEG data were re-referenced by subtracting the average across all sensors excluding those marked as bad. This was only performed for the LEMON dataset. This was the last step in preprocessing the sensor-level EEG data.
6. **Coregistration.** Scalp, inner skull, and outer brain surface were extracted from the sMRI using FSL BET (Smith, 2002; Jenkinson, 2005). The nose was not included in the scalp surface because the sMRI images were defaced. The scalp surface was then used in OSL RHINO (Quinn et al., 2022) to coregister the sMRI to the M/EEG data based on matching the scalp to a set of digitised Polhemus head shape points and fiducials.
7. **Forward modelling.** The *forward model* gives us the expected signal at each sensor (leadfield) for a dipole placed at a particular location inside the brain. The electric/magnetic field from this dipole transverses different tissue types, which have different electromagnetic properties. It is common in MEG source localisation to model the changes in magnetic properties using a single boundary separating the volume inside the brain from outside. In the current work, we used a boundary element model (Henson et al., 2009), using the inner skull surface as the boundary. For EEG, we need to model the electrical conductivity for the different tissue types and used three layers in the forward model: the inner skull, outer skull and outer head skin. We calculated the leadfields for a dipole placed in the *x*, *y* and *z*-direction on an 8 mm isotropic grid inside the volume bound by the inner skull.
8. **Bandpass filtering and bad segment removal.** We applied a bandpass filter to the preprocessed M/EEG data (fifth-order IIR Butterworth) to focus on the frequency range of interest. Depending on the dataset we used a different frequency range: 0.5-80 Hz for Cam-CAN; 1-45 Hz for BioFIND, MEGUK and LEMON. Following this, we removed bad segments from the data to ensure these did not impact on the quality of the source localisation.
9. **Volumetric beamforming.** We employed a beamformer (Van Veen and Buckley, 1988; Hillebrand and Barnes, 2005; Sekihara and Nagarajan, 2008) to source localise the MEG data. The beamformer estimates source activity (at a location inside the brain) as a linear combination of sensor activity. In this step, we calculated the weights for the linear combination, referred to as the *beamformer weights*. We used a unit-noise-gain invariant Linearly Constrained Minimum Variance (LCMV) beamformer. This type of beamformer controls for the depth bias by normalising the beamformer weights (Tait et al., 2021). To calculate the beamformer weights, we need to provide the leadfields from the forward model, the covariance of noise (signal measured by the sensors unrelated to brain activity) and the covariance of the data (signal measured by the sensors containing both noise and brain activity). For the noise covariance, we used a common choice for when empty room recordings are not available: a diagonal matrix containing the variance of each sensor type. The diagonal values are chosen to account for the scaling of each sensor type. We filled the diagonal elements with the average (temporal) variance across the sensors of that type. For the data covariance, we used the covariance of the data from step 8 reduced to a rank of 60 for the MEG datasets and 54 for the EEG dataset (this regularisation improves the robustness of the beamformer). The rank was chosen to be less than the full rank of the data (∼64 for MEG and ∼61 for EEG) and above the number of parcels (52; see step 10). The beamformer calculates the activity in the *x, y* and *z*-direction at each grid point used in the leadfield calculation. This is reduced to a single value by projecting the *x, y, z* activity onto the axis that maximises power of at that location. After beamforming, we are left with the activity at each grid point inside the brain, referred to as a *voxel*.
10. **Parcellation.** Often, we have a very large number of voxels, so we parcellate the data into particular regions of interest (ROIs). In the current work, we used an anatomically defined 52-ROI parcellation. See (Kohl et al., 2023) for a description of the parcellation. Parcel time courses were obtained by applying principal component analysis (PCA) to the (demeaned) voxel time courses assigned to a parcel and taking the first principal component. PCA is preferred over simply taking the mean across voxels because there is a sign ambiguity in the voxel time courses due to the beamformer.
11. **Orthogonalisation.** The source localisation of M/EEG data suffers from spatial leakage where the uncertainty in estimating source activity leads to nearby locations in source space exhibiting highly correlated activity. These correlations can be misinterpreted as functional connectivity (so called ‘ghost interactions’ (Colclough et al., 2015)). To minimise the impact of spatial leakage, we used the symmetric orthogonalisation technique proposed in (Colclough et al., 2015), which removes all zero-lag correlations from the parcel data. This can be seen as a conservative approach as we have likely removed some genuine zero-lag functional connectivity from the data.
12. **Sign flipping.** Unfortunately, there is an ambiguity in the sign of each parcel time course (due to the ambiguity of the dipole direction in the source reconstruction and due to the PCA used in the parcellation). This means the sign of each parcel time course can be misaligned across subjects. After we have the orthogonalised parcel time courses for each subject, we calculate the covariance of time-delay embedded parcel data (see Section 2.3.1) for each subject and flatten the upper triangle into a vector. Comparing the correlation of this vector for each pair of subjects, we selected the median subject (highest average correlation) as a template. Finally, we match the sign of each subject’s orthogonalised parcel time course to the template with the random search sign-flip algorithm described in (Vidaurre et al., 2016) using the correlation between flattened time-delay embedded covariance matrices as the metric for alignment.
13. **Standardisation.** Finally, we standardise the data by subtracting the mean and dividing by the variance for each parcel individually.

Note, in the current work all references to the ‘parcel time courses’ are to the standardised, sign-flipped, orthogonalised parcel time courses.

### 2.3 Parcel-Level Canonical HMM

Here, we describe how we developed a parcel-level canonical HMM based on the Cam-CAN dataset.

#### 2.3.1 Data Preparation

Before training the HMM on Cam-CAN, the parcel time courses were prepared by performing time-delay embedding (TDE), which augments the data with extra channels corresponding to time-lagged versions of the original data. The rationale for this is to encode spectral information (frequency of oscillations) into the covariance matrix of the data. This is discussed further in section 5.4 of (Gohil et al., 2024a). In the current work, we used ±7 lags (±28 ms), resulting in 780 channels for the TDE data. PCA was then applied at the group level to reduce the TDE data down to 120 channels. Finally, we (temporally) standardised (demeaned and divided by the standard deviation) the TDE-PCA data. These were the data used to train the HMM. These steps are summarised in Figure S2.

#### 2.3.2 Model Training

The HMM (Rabiner and Juang, 2003; Bishop and Nasrabadi, 2006) is a generative model typically used to study time series data. At each time point, *t*, there is an underlying categorical hidden state, *θ_t_* ∈ {1, …, *K*},where *K* is the number of states. The observed time series, ***x****_t_*,is generated using a multivariate Normal distribution based on the hidden state:

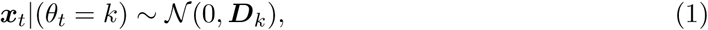

where *θ_t_* = *k* is the state at time point *t* and ***D****_k_* is a *state covariance*. Note, we force the mean to be zero to focus on modelling dynamics in the covariance, which contains the connectivity information we are interested in. Dynamics are governed by transitions in the hidden state. Note, it is assumed that the probability of a transition at time point *t* only depends on the state at the previous time point *θ_t–_*_1_ (this is the Markovian constraint).^2^ Each pairwise state transition probability is contained in the *transition probability matrix*:

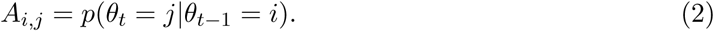

When we train an HMM on data, we learn the most likely value for the state covariances, {***D***_1_, …, ***D****_K_* }, and transition probability matrix, ***A***, to have generated the data. We use stochastic variational Bayes (Bishop and Nasrabadi, 2006) to do this, where we iteratively update our estimates for {***D***_1_, …, ***D****_K_*, ***A***} based on a random subset of the training data to minimise the variational free energy. We use the Baum-Welch algorithm (Bishop and Nasrabadi, 2006) to calculate the (*posterior*) probability of each state being active at each time point, *q*(*θ_t_*), for each session based on our estimates for {***D***_1_, …, ***D****_K_*, ***A***}. The posterior, *q*(*θ_t_*), is a (states, time) array for each session. The training of an HMM is summarised in Figure S3.

##### Hyperparameters

We used the HMM implemented in the osl-dynamics (Gohil et al., 2024a) toolbox. In order to train an HMM, we need to specify a few hyperparameters. An important hyperparameter is the number of states. In the current work, we trained canonical HMMs with 4-16 states, which is in line with previous HMM M/EEG studies (Vidaurre et al., 2018b; Cho et al., 2024; Gohil et al., 2024a; Guichet et al., 2025; Rossi et al., 2023; Fauchon et al., 2022; Higgins et al., 2021; Kohl et al., 2025; Seedat et al., 2023; Sitnikova et al., 2018; Gohil et al., 2024b; van Es et al., 2023; Gohil et al., 2022; Quinn et al., 2018; Vidaurre et al., 2018a, 2016; Coquelet et al., 2022; Zhang et al., 2021). Other hyperparameters include the batch size, sequence length and learning rates. These had little impact on the HMM inference. The values used are summarised in Table S1.

##### Run-to-run variability

The final estimates for {***D***_1_, …, ***D****_K_*, ***A***} can be sensitive to the initial values used at the start of training. The typical approach for overcoming this is to train several models from scratch starting from random initialisation and picking the one with the lowest final variational free energy (deemed the best description of the data) for the subsequence analysis. Historically, this has produced very reproducible results (Gohil et al., 2024a). In our case, we analyse a particularly large dataset. This makes the HMM inference very stable, and we consistently converge on very similar estimates for {***D***_1_, …, ***D****_K_*, ***A***}. Despite this, we took a cautionary approach and selected the best model from a set of five runs.

### 2.4 Applying the Parcel-Level Canonical HMM to a Boutique Dataset

#### 2.4.1 Data Preparation

Before we can apply the canonical HMM to a new boutique dataset, we must prepare the parcel time courses in a similar way to how the Cam-CAN data was prepared when training the canonical HMM, i.e. using the steps described in Section 2.2. Importantly, we use the same parcellation and a matching sampling frequency (250 Hz). We also strongly recommend using the template session from the Cam-CAN dataset when doing sign flipping on the new dataset, and using the PCA components from Cam-CAN when applying PCA following TDE.

#### 2.4.2 Inference (Estimating State Probabilities)

We use the canonical HMM pre-trained on Cam-CAN to perform inference on a new dataset. This is shown in Figure S4. In doing this, we obtain a state probability time course for each session in the new dataset.

#### 2.4.3 Dual Estimation: Individualised Post-Hoc Analysis

We use the state probability time course and parcel time course for each session to perform post-hoc analysis, see Figure S5. This process provides session-specific estimates for network properties and is referred to as *dual estimation* (Vidaurre et al., 2017).

##### State power/coherence spectra

Here, we combine the state probability time course with the original parcel data (pre-TDE-PCA) to estimate the spectral properties (PSD and coherence) of each state using a multitaper (Babadi and Brown, 2014). This involves performing the following steps for each session and state:

1. Temporally standardise (demean and divide by the standard deviation) the parcel time courses and multiply by the state probability time course.
2. Calculate power/cross spectral density (P/CSD) for each parcel/pair of parcels using a multitaper (2 s window, 0% overlap, 7 DPSS tapers, 4 Hz time half bandwidth). This is the standard approach/settings used for HMMs trained on TDE-PCA data (Gohil et al., 2024a; Vidaurre et al., 2018b, 2016).
3. Use 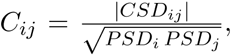 where *i* and *j* correspond to parcels, to calculate the coherence from the P/CSD.
4. The amplitude of the PSD is biased by the amount of time each state is active due to the multiplication in step 1. We account for this by scaling the PSD by one over the fractional occupancy of the state.

##### State power maps

The multitaper results in a [sessions, states, parcels, frequencies] array containing the PSDs. Averaging over the frequency dimension results in a [sessions, states, parcels] array, which we refer to as the *state power map* for each session. In the visualisation of the group-averaged power maps, we displayed each state’s power map separately and used the average (parcel-specific) power across states as the reference.

##### State coherence networks

The multitaper results in a [sessions, states, parcels, parcels, frequencies] array for the coherences. Similar to the state power maps, we reduce the frequency dimension by taking the average over the frequency dimension. This results in a [sessions, states, parcels, parcels] array, which represents the *state coherence network* for each session. We displayed the group-average coherence network for each state individually using the (edge-specific) average across states as the reference and thresholding the top 3% of edges irrespective of sign.

##### Summary statistics for dynamics

We calculated a *state time course* by taking the most probable state (*maximum a posteriori probability estimate*) at each time point. From the state time course, we calculate metrics that summarise the dynamics within a session. For each session and state, we calculated:

- Fractional occupancy: the fraction of total time spent in a state.
- Mean lifetime (ms): the average duration a state was active.
- Mean interval (s): the average duration between successive state activations.
- Switching rate (Hz): the average number of state activations per second.

### 2.5 Sensor-Level Canonical HMM

Alongside the parcel-level canonical HMM, we developed a corresponding sensor-level version. Importantly, the sensor-level canonical HMM was designed to capture the same states as the parcel-level canonical HMM. This allows researchers to apply the sensor-level canonical HMM to new sensor-space datasets without the need for source reconstruction, while still being able to map to the same set of canonical networks. Here, we describe how the sensor-level canonical HMM is obtained (i.e. trained) and how to apply it to new sensor-space datasets.

#### 2.5.1 Data Preparation

The sensor-level canonical HMMs were inferred using the MaxFiltered Cam-CAN data (Elekta, *N* = 621, rest and task). We first bandpass filtered the data between 1 and 45 Hz to remove noise. Due to the MaxFiltering the rank of the sensor-level data was reduced to 64. Therefore, after filtering we use PCA to reduce to 64 channels. The data was then prepared in the same way as the parcel time courses with TDE-PCA using ±7 lags followed by a second (post TDE) PCA. The number of components used in the second PCA was chosen to maximise the agreement (classification accuracy) between the sensor and parcel-level inferred state time courses. The optimum number of PCA components was found to be 160.

#### 2.5.2 Model Training

We seek a canonical HMM that captures the same states as those obtained by an HMM trained on the (prepared) parcel data directly. Therefore, to calculate the sensor-level canonical HMM we did not train a new model from scratch. Instead, we dual estimated the state covariances using the prepared *sensor-space* Cam-CAN data and the state probability time courses inferred using the parcel-level canonical HMM. In the dual estimation, the states were assumed to have zero mean and we used the transition probability matrix/initial state probabilities from the *parcel-level* canonical HMM. The result of this was group-level state canonical covariances for each HMM state in prepared sensor-space.

#### 2.5.3 Application to Boutique Datasets

The sensor-level canonical HMM can be applied to boutique datasets in the same way as the parcel-level HMM. The only difference is that the input data must be the prepared sensor-level data and must have the same sensor layout as in the canonical HMM. Currently, the public repository for code (see Section 6) provides sensor-level canonical HMMs for Elekta MEG data.

### 2.6 Modelling Approaches: Conventional vs Canonical HMM

Two different modelling approaches were used in this work: a conventional approach, where a new HMM is trained from scratch on the boutique dataset; and a canonical approach, where a pre-trained HMM is used. No HMM training is required in the canonical approach. An overview of each approach is shown in Figure 1. Both approaches use the same preparation steps on the new dataset and the same dual estimation step to obtain individualised networks for each session. Note that in all results we show using the conventional approach, the run with the lowest final variational free energy from a set of 10 was used.

**Figure 1:**
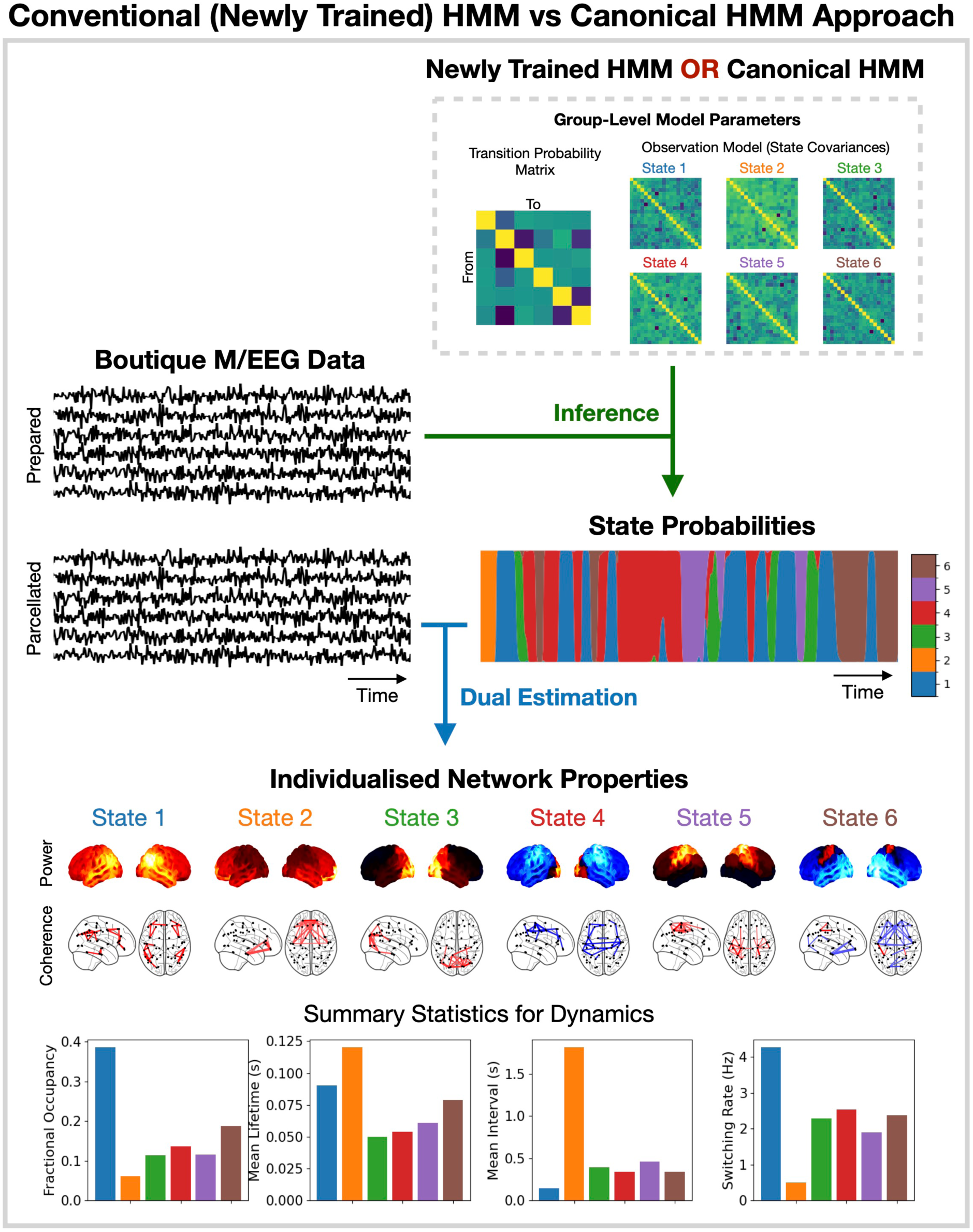
Overview of HMM approaches. A conventional HMM approach involves training a new model on the boutique dataset (usually multiple times to ensure results are robust), whereas the canonical HMM approach uses a HMM pre-trained on a large independent dataset. The HMM is used to infer the state probabilities at each time point and dual estimation is used to infer individualised network properties for each state.

## 3 Results

We present a canonical HMM that describes brain activity using a set of transient networks derived from the Cam-CAN dataset. We describe the spatio-spectral patterns of brain activity associated with each network (in parcel space) and summarise each network’s dynamics in Section 3.1. Applying the parcel-level canonical HMM to a boutique MEG dataset in Section 3.2, we study how mild cognitive impairment, an early indicator of Alzheimer’s disease, alters brain network activity. In Section 3.4, we illustrate how a parcel-level canonical HMM from MEG can be applied to parcellated EEG data. Finally, we present a sensor-level canonical HMM that can be applied to new sensor-level MEG data in Section 3.5; allowing for a more flexible application of the canonical HMM, where the user does not need to apply the same source localisation used in the parcel-level model.

### 3.1 Parcel-Level Canonical HMM

We start by presenting the network description provided by the parcel-level canonical HMM for a model order (number of states) of 10. Note that canonical HMMs for a range of model orders (4 to 16) have been made available on the code repository (see Section 6). The 10state canonical HMM is shown in Figure 2. The canonical HMM was trained on 1849 MEG recording sessions (rest and task) from the Cam-CAN dataset. Each HMM state corresponds to a network of brain activity with fast transient dynamics (∼ 100 ms). These networks describe recurrent spatio-spectral patterns found in a large population of healthy individuals across a wide age range (*N* = 621, 18-88 years old). Figure 2 shows the canonical HMM networks at the group level (i.e., group-averaged). Individualised networks (for different sessions) are shown in Figure S6. Figure 2 shows the network activity in the frequency range 1-45 Hz, which is the typical frequency range used in previous studies (Gohil et al., 2024a). However, the canonical HMM was trained on 0.5-80 Hz (prepared) parcel data (see Section 2.2 for the full details). This was done to ensure the canonical HMM can be applied to the widest frequency range possible discussed further in Section 4.4.

**Figure 2:**
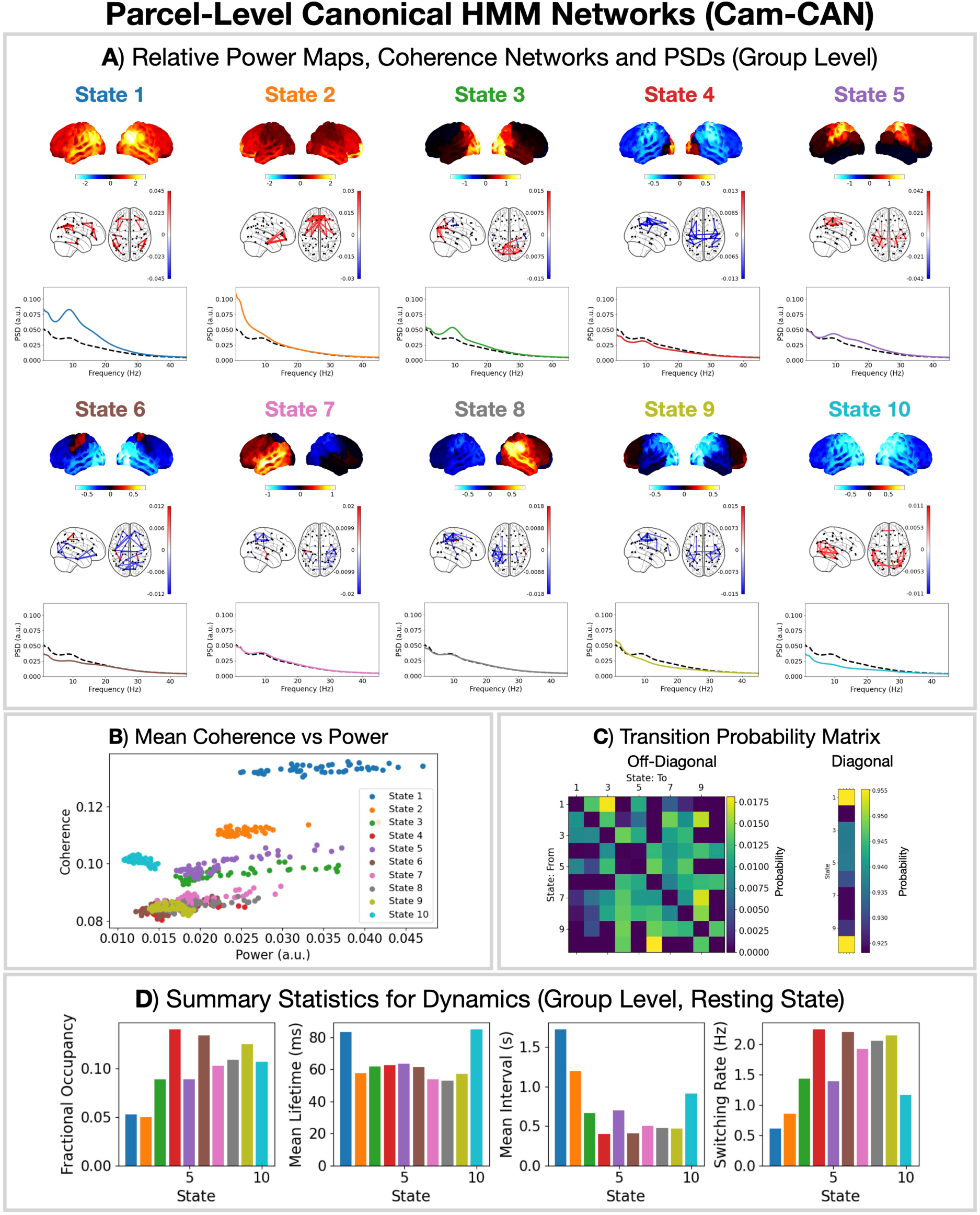
10 state parcel-level canonical HMM networks inferred from Cam-CAN MEG data (*N*=621, rest and task). A) Relative state power maps, coherence networks (top 3%) and PSDs showing activity in the frequency range 1-45 Hz. The solid line shows the state PSD averaged over regions (parcels) and the dashed line shows the PSD averaged across states. B) Mean coherence vs power for each parcel averaged over subjects. C) State transition probability matrix. D) State summary statistics for dynamics: fractional occupancy; mean lifetime; mean interval; and switching rate.

### 3.2 Example MEG Boutique Study: Resting State

Here, we illustrate the use of a 10 state parcel-level canonical HMM in studying a boutique (i.e. bespoke) resting-state MEG dataset. We adopted a case-control analysis (comparing the networks for healthy vs. diseased groups) using the BioFIND dataset (Hughes et al., 2019; Vaghari et al., 2022), which aims to identify biomarkers for the early detection of Alzheimer’s disease (Scheltens et al., 2021). This study collected MEG data (resting state, eyes closed) from individuals at high risk of developing Alzheimer’s disease. These participants showed early signs of cognitive decline, referred to as *Mild Cognitive Impairment* (MCI). See Section 2.1.2 for details regarding the dataset.

The conventional approach for a case-control HMM analysis would be to first train a new HMM on the full BioFIND dataset by concatenating the data from all participants (healthy and MCI) (Gohil et al., 2024a). The HMM is unaware of which participant the data comes from (i.e. it is unsupervised) and finds the best set of network states that describes the entire dataset. This model is referred to as a *group-level HMM*. Following this, subject-level networks are estimated for each participant post-hoc based on the group-level state activations using *dual estimation* (Vidaurre et al., 2017), see Figure S5. By first training the group-level HMM, we ensure the networks are common to all individuals/groups, i.e. state 1 for one individual is the same network as state 1 for another. Figure 3A shows the group-level HMM networks adopting a conventional approach to BioFIND. This shows a set of networks similar to those typically produced from an HMM analysis (Gohil et al., 2024a).

**Figure 3:**
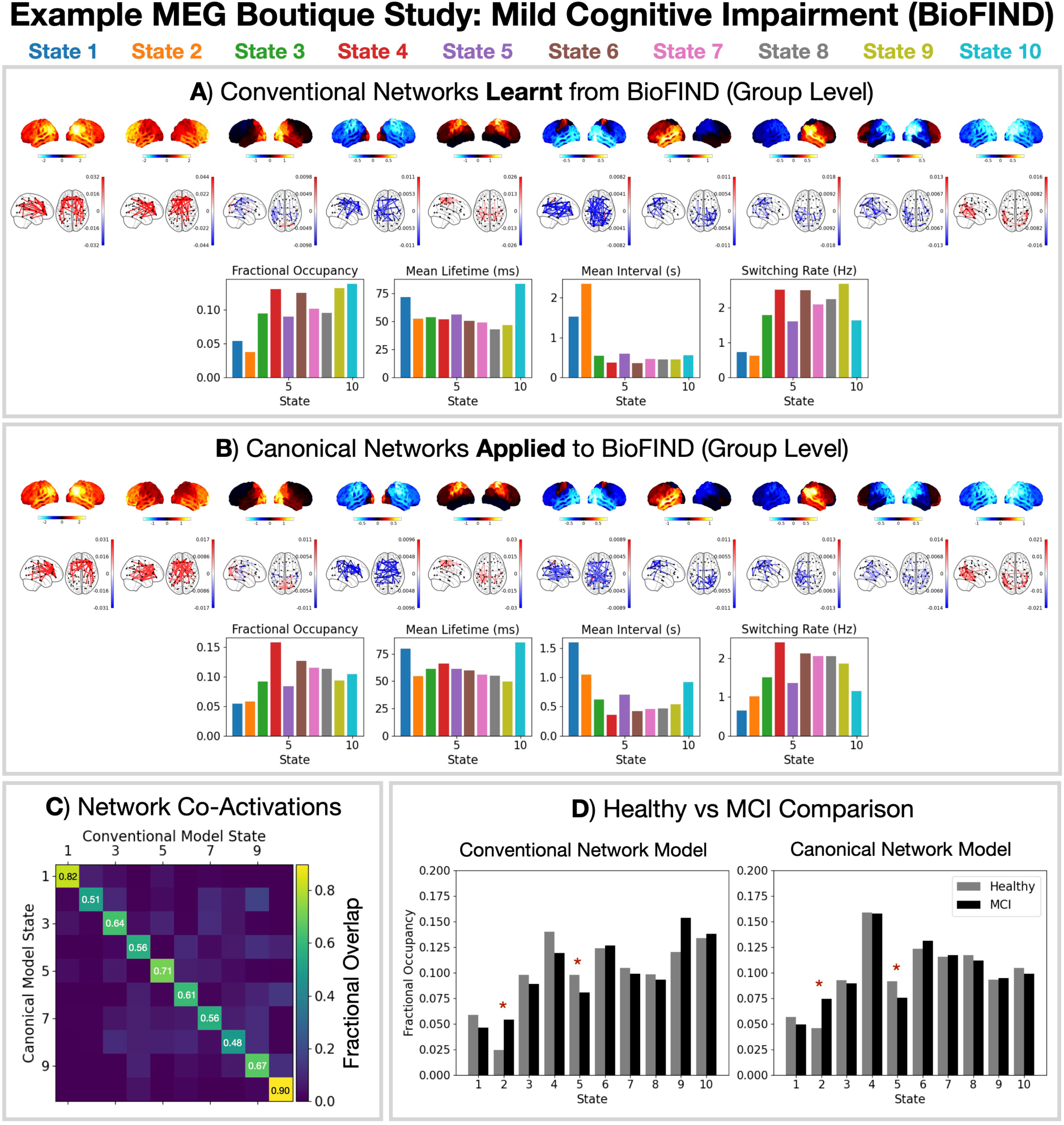
Applying the canonical HMM approach provides a common set of networks that can be compared across studies and reduces the computational resources needed. A conventional (A) and canonical (B) HMM modelling approach was applied to the BioFIND dataset. Relative state power maps (1-45 Hz), coherence networks (1-45 Hz, top 3%) and summary statistics for dynamics are shown. State activations from each approach show good agreement (normalised confusion matrix shown in C) and the same differences between mild cognitive impairment (MCI) and healthy control groups (D). Group differences were calculated using a General Linear Model (GLM) with age and sex as confounds. Row-shuffle permutations were used for statistical significance testing with the maximum *t*-statistic across all features and states being used to control for multiple comparisons. The asterisk (*) indicates a *p*-value < 0.05.

With the canonical HMM approach, we do not need to train a new HMM on the BioFIND dataset. We can instead use the *pre-trained* canonical HMM at the desired model order. We just need to ensure that BioFIND has been processed into the same parcellated source space as the canonical HMM. We can infer the state probabilities using the canonical HMM and proceed to the dual estimation. Figure 3B shows the group-level HMM networks from applying the canonical HMM to BioFIND. Qualitatively, we see the same networks as the conventional HMM approach (Figure 3A). However, there are two key benefits of adopting the canonical HMM approach: first, it saves substantial computation by avoiding the need to train a new model; second, it provides a common reference for comparing different studies and datasets.

A quantitative comparison of the agreement of the inferred states using the conventional and canonical HMM is shown in Figure 3C. We see both approaches have a high agreement in the state that is assigned at each time point (0.5-0.9 fractional overlap; chance = 0.1). Turning to the case-control analysis, we see in Figure 3D, we see that both approaches give the same significant differences in the fractional occupancy of the states. MCI patients increase the time spent in state 2 (the frontal network) and decrease the time spent in state 5 (the sensorimotor network).

### 3.3 Example MEG Boutique Study: Task

Next, we illustrate the use of the 10 state parcel-level canonical HMM on a boutique task MEG dataset (MEGUK Partnership, 2025). The MEGUK dataset was recorded while the participant performed a visual *N*-back task with 0, 1, and 2-back conditions. This task involves higher-order cognitive processes such as working memory (Owen et al., 2005). We applied a conventional HMM network response analysis (described in (Gohil et al., 2024b)) to the MEGUK dataset and compare it to the canonical HMM.

Figures 4A and B show we infer qualitatively similar networks are inferred using the conventional and canonical HMM approach. Figure 4C quantifies the agreement, we see a fractional overlap of 0.2-0.9 (chance = 0.1) for the two approaches.

**Figure 4:**
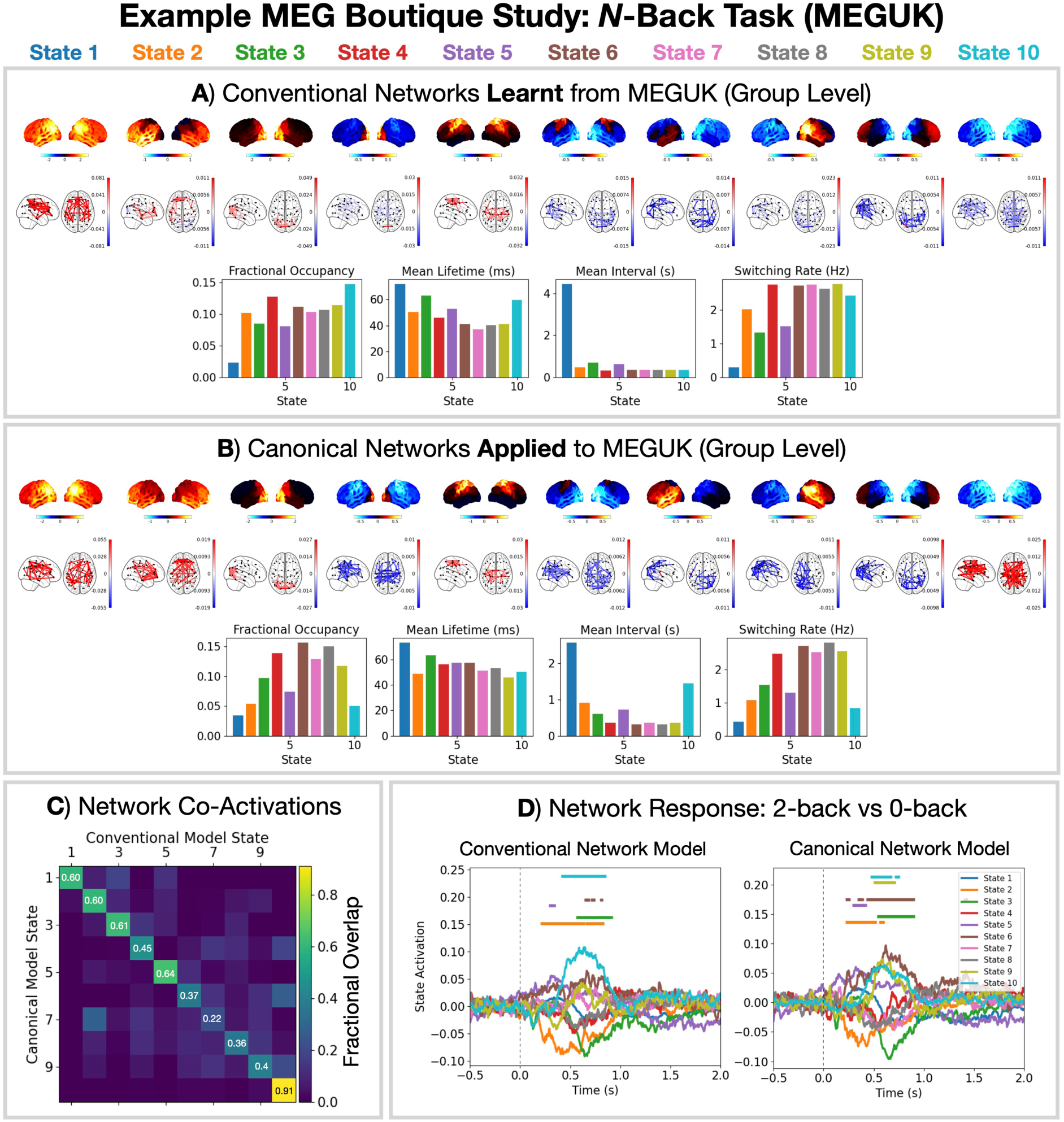
The canonical HMM networks can describe brain activity related to higher-order cognitive processes. A conventional (A) and canonical (B) HMM modelling approach was applied to the MEGUK dataset. Relative power maps (1-45 Hz), coherence networks (145 Hz, top 3%), and summary statistics for dynamics are shown. State activations from each approach show good agreement (normalised confusion matrix shown in C) and the same network response for the 2-back minus 0-back condition is observed (D). The dashed vertical line is the presentation of the stimulus. Horizontal bars indicate time points with *p*-value < 0.05, which were calculated using sign-flip GLM permutations taking the maximum statistic (COPE) across time and states to control for multiple comparisons.

Figure 4D shows the baseline corrected network response: the state time course epoched around the presentation of visual stimuli and averaged over trials and subjects (Gohil et al., 2024b). We took the difference between the trial-average for the 2-back condition and 0-back condition before calculating the subject average. This gives the network activity related to working memory while controlling for visual processing. We see an early deactivation of state 2(*t* = 0.1-0.5 s) followed by an activation of state 9 and 10 (*t* = 0.5-1 s). Additionally, we see an activation of states 5 and 6. Both the conventional and canonical HMM approach show good agreement, with the canonical HMM identifying more significant network responses. The network responses presented here generally agree with a conventional HMM approach applied to an independent *N*-back task dataset in (Rossi et al., 2023). The functional relevance of the canonical HMM networks is discussed in Sections 4.8 and 4.9.

### 3.4 Example EEG Boutique Study

The canonical HMM was inferred on parcellated MEG data (see Section 2). However, source activity within the brain can be estimated using other neuroimaging modalities. EEG offers a cheap non-invasive measure of brain activity with high temporal resolution, albeit at some cost to the spatial resolution and signal-to-noise ratio (Blinowska and Durka, 2006). The same source reconstruction methods as MEG (see Section 2.2) can be applied to medium/high-density (64-256 electrode) EEG recordings (Cho et al., 2024; Makkinayeri et al., 2025). Once the EEG data has been source reconstructed into the same parcel space, we can apply the canonical HMM approach to identify MEG-like networks using EEG. Here, we use the LEMON dataset and compare a conventional HMM approach with the canonical HMM approach.

The conventional HMM approach, i.e. training a new HMM from scratch on the (prepared) parcellated EEG data, is studied in (Cho et al., 2024). Figure 5A shows the spectral properties and dynamics this produces. We see training a new HMM on parcellated EEG data identifies similar networks to parcellated MEG, see (Cho et al., 2024) for a quantitative comparison. However, a few networks that are often identified in MEG are not found: the sensorimotor networks and the temporal networks (states 6, 7, 8 in Figure 2).

**Figure 5:**
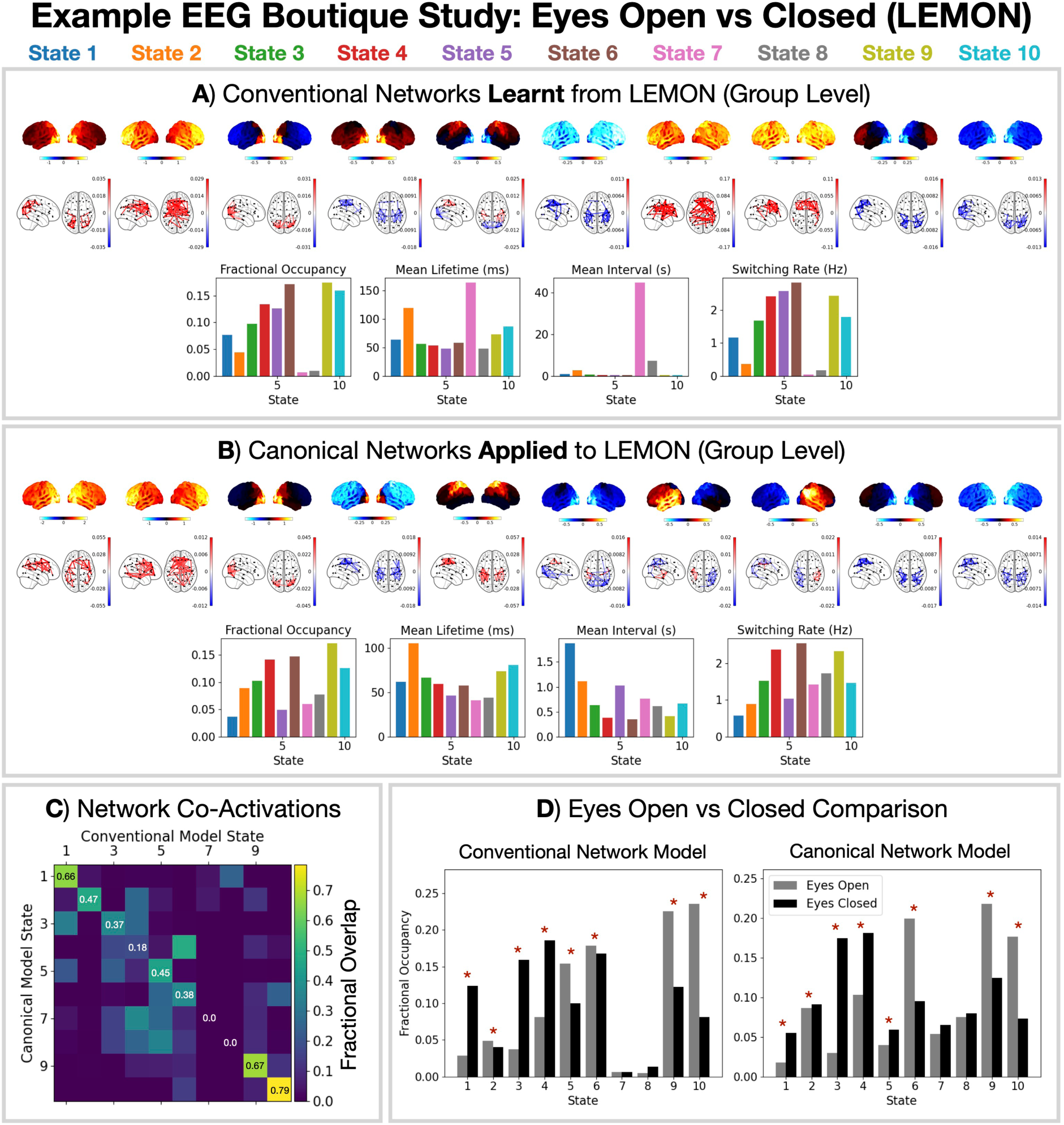
MEG-like networks can be inferred from parcellated EEG data. A conventional (A) and canonical (B) HMM modelling approach was applied to the BioFIND dataset. Relative power maps (1-45 Hz), coherence networks (1-45 Hz, top 3%), and summary statistics for dynamics are shown. Fractional verlap between state activations from each modelling approach (i.e. normalised confusion matrix) is shown in C. The same differences in state fractional occupancy for an eyes open vs closed contrast are found in both modelling approaches (D). Contrasts were calculated using a GLM with row-shuffle permutations for statistical significance testing using the maximum *t*-statistic across states to control for multiple comparisons. The asterisk (*) indicates a *p*-value < 0.05.

When we adopt the canonical HMM approach, we can identify the states missing from the conventional analysis (Figure 5B). Figure 5C shows the agreement between the state time courses from each modelling approach. We see states 7 and 8 from the conventional model do not correspond very well with any of the canonical HMM networks, suggesting they represent dataset-specific features or noise. The resting-state recordings in the LEMON dataset alternated between 30 second periods of eyes open and closed conditions. In Figure 5D, we contrast the fractional occupancy of each state in each condition. We see generally good agreement in the change in fractional occupancy for the two modelling approaches.

### 3.5 Sensor-Level MEG Canonical HMM

Applying a parcel-level canonical HMM to a boutique dataset requires the data to have been processed in the same way as the training data for the canonical model (including source reconstruction and parcellation). Example scripts for how to do this have been provided, see Section 6. A useful alternative approach would be to apply the HMM at the sensor-level, thus avoiding the need for source reconstruction. This facilitates broader applications in scenarios where source reconstruction is not feasible, such as in real-time network detection. It was recently shown in EEG that a reasonable agreement in state classification can be achieved between HMMs inferred at the sensor and parcel level (Cooray et al., 2024).

We developed a canonical HMM that can be applied to sensor-level MEG data, albeit with the requirement that the data have the same sensor layout as the Cam-CAN dataset, i.e. correspond to an Elekta MEG scanner. Details of how the sensor-level canonical HMM was derived are provided in Section 2.5. Notably, the sensor-level canonical HMM was designed to capture the same states as the parcel-level canonical HMM. As with the source-level canonical HMMs, we provide sensor-level canonical HMMs for different model orders (4-16 states).

Figure 6A shows the relative power maps and PSDs for the 10-state sensor-level canonical HMM at the sensor level. Qualitatively, we see a good correspondence between the sensor-level and parcel-level power maps. Note that, by virtue of the sensor-level canonical HMM being obtained via the parcel-level canonical HMM, the states are naturally in the same order.

**Figure 6:**
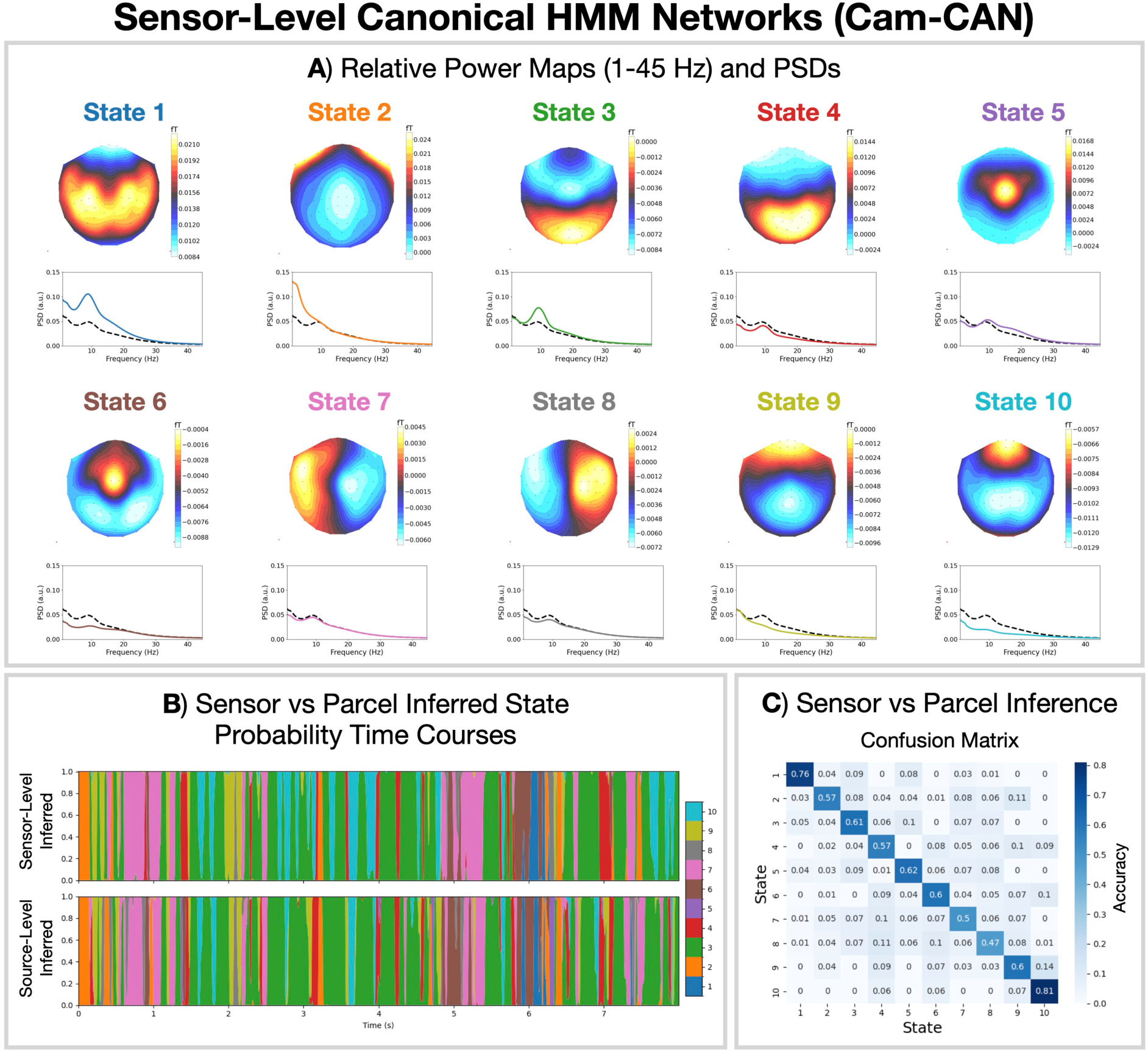
The canonical HMM approach can be applied in MEG sensor space. This allows networks to be approximately inferred when source reconstruction is not feasible. A) Topographic maps of the relative state power (1-45 Hz) at each MEG sensors viewed from above and PSDs for the 10-state canonical HMM networks at the sensor level, in the same order of states as for the parcel-space canonical HMM shown in Figure 2. The solid line shows the state PSD averaged over regions (parcels) and the dashed line shows the PSD averaged across states. B) Illustrative state probabilities inferred from the first 8 seconds of the first session in Cam-CAN when applying the canonical HMM networks to the sensor-level data (top) or the (prepared) parcellated data (bottom). C) Normalised confusion matrix: average accuracy (across participants) for the state assignment at each time point between the sensor-level and source-level canonical models.

Figure 6B shows the inferred state probabilities when applying the canonical HMMs at the sensor (top) and parcel (bottom) levels. Quantifying the agreement using a confusion matrix (which counts the number of correct and incorrect classifications between the sensor and source-level state assignment) in Figure 6C, we see the inferred states agree very well. In particular, states 1 and 10 can be identified with a high accuracy at the sensor level: approximately 0.8 vs. a chance accuracy of 0.1.

## 4 Discussion

We propose a canonical HMM that can be used to study transient networks in M/EEG data. Each HMM network corresponds to a unique spatio-spectral pattern of brain activity with fast dynamics (Figure 2). The canonical HMM was trained on rest and task (visual, auditory, sensorimotor) data from a large cohort of healthy participants (*N* = 621, 18-88 years old, Cam-CAN), capturing a diverse range of brain activity and population variability. We demonstrated how the parcel-level canonical HMM could be used to analyse a boutique resting-state MEG dataset for early Alzheimer’s detection (Figure 3) and task MEG dataset for studying working memory (Figure 4). We showed the canonical HMM networks could be transferred to parcellated EEG data (Figure 5). We also provided a sensor-level canonical HMM for Elekta MEG (Figure 6). Here, we discuss some considerations regarding the canonical HMM and highlight some advantages of adopting this approach.

### 4.1 Canonical HMM Networks as a Basis Set

The canonical HMM states presented in this work can serve as a basis set for studying M/EEG data using a set of canonical transient networks. These networks were inferred on a large MEG cohort, which is sufcient to ensure good reproducibility on unseen datasets (Figures 3 and 5). A benefit of taking this approach is it provides a description of the data in a common space, which can be useful in comparing individuals, especially in boutique or individualised (*N* = 1) studies. The adoption of a common basis set also improves the comparability of analysis across different studies. Conventional M/EEG HMM studies typically involve training a model from scratch. This means the networks inferred in one study do not exactly match another. This can be resolved via the adoption of a common canonical HMM.

### 4.2 Reduction in Computational Resources Needed

A key benefit of adopting a canonical HMM approach is it means the computational resources needed to perform an HMM analysis have been greatly reduced. The end user no longer needs to train their own HMM, which can be a computationally demanding process that often needs to be repeated many times. The canonical HMMs trained in this work utilised modern GPU hardware and took on the order of 10 hours to complete a single run. In contrast, applying a trained HMM to new data (inference) requires relatively little computational resources and is quick (on the order of minutes). HMM inference can be done straightforwardly on most modern CPUs. This increases the accessibility of these tools to end users without access to significant computational resources.

### 4.3 Stable Network Inference

As with many machine learning approaches, the HMM can suffer from a local optima issue, where the model can converge to different final solutions (model parameters). Conventionally, this is resolved by training many HMMs from scratch and selecting the best for subsequent analysis (usually the one with the lowest variational free energy (Gohil et al., 2024a)). Adopting a canonical HMM alleviates this issue because given the trained model and data, the inference is deterministic.

### 4.4 Preprocessing and Source Reconstruction

To apply the parcel-level canonical HMM to new data, it must be preprocessed and source reconstructed in a similar manner to the data used to train the canonical HMM. This is one of the main limitations of the proposed method. The key data processing steps that need to be matched are downsampling to 250 Hz, LCMV beamforming, parcellation with the same atlas, orthogonalisation, sign flipping to match the template subject from Cam-CAN, number of time-delay embeddings, and PCA using the same components from Cam-CAN. All the necessary data and example scripts for this process are provided in the code repository (see Section 6). Other steps such as the exact band-pass filtering range and data cleaning are less critical.

Figure 7 shows the agreement in state classification for data processed with a different band-pass filter and parcellation. Previous HMM studies have focused on the frequency range 1-45 Hz (Gohil et al., 2024a), which is why we used a 1-45 Hz bandpass filter in the boutique studies. In this work, we provide a canonical HMM trained for a wider frequency range: 0.5-80 Hz. This was done to ensure the canonical HMM captures dynamics across the widest possible frequency range of interest. Applying the canonical HMM to data bandpass filtered over a narrower range (1-45 Hz) yields very similar state time courses to the full range (0.5-80 Hz; fractional overlap of 0.75-0.86; Figure 7A). Using a different parcellation shows a lower fractional overlap between state time courses (0.41-0.78; Figure 7B), however, there is still a good one-to-one correspondence in state assignment.

**Figure 7:**
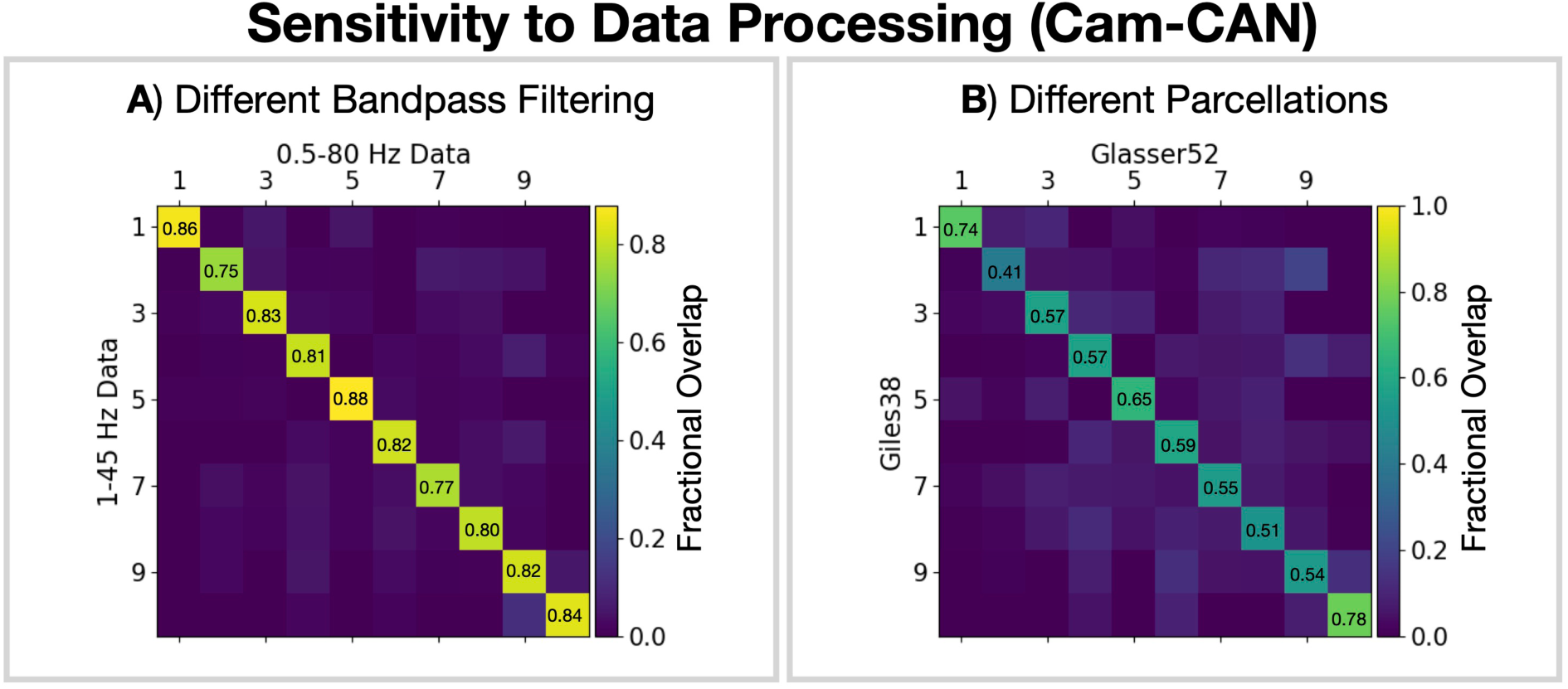
The canonical HMM infers consistent state time courses for differing data processing choices. Fraction of time points where state assignments agree (i.e. normalised confusion matrix) for data bandpass filtered over different frequency bands using the Glasser52 parcellation (A) and different parcellations using a 0.5-80 Hz bandpass filter (B). State time courses were inferred on the full Cam-CAN dataset.

For scenarios where the end user does not wish to adopt the same source reconstruction, we provide a sensor-level canonical HMM for Elekta MEG.

### 4.5 Transferring Elekta MEG Networks to Other Modalities

The objective of this work was to develop a canonical HMM based on MEG. In this work, we used Elekta MEG data to train the canonical HMM. However, this model can be leveraged in studies using other modalities/scanners if we are able to source reconstruct and parcellate the data. This is discussed further below.

#### 4.5.1 Different MEG Scanners and OPMs

In this work, we applied the canonical HMM to boutique MEG datasets collected using an Elekta scanner (Figure 3) and CTF scanner (Figure 4). Elekta and CTF scanners are based on Superconducting Quantum Interference Device (SQUID) technology for measuring the brain’s magnetic field (Hämäläinen et al., 1993). Although these scanners are based on the same technology, they can have different sensitivities and noise profiles (Hämäläinen et al., 1993), which means accounting for scanner differences is often important. In this work, we source reconstruct and parcellate the data, which acts as a powerful harmonisation step that can minimise these differences. Consequently, a canonical HMM trained on Elekta data can transfer well to CTF data as was the case in Figure 4. However, it is certainly possible that systematic biases related to the scanner remain, which will affect the canonical HMM state inference. Therefore, the user should note the (current) canonical HMM was trained on Elekta data only and should control for scanner differences in subsequent analysis.

Recently, a new device for recording the MEG signal has been developed: Optically Pumped Magnetometers (OPMs) (Boto et al., 2018). The consistency of SQUID-based recordings and OPM recordings after source reconstruction was demonstrated in (Tanner et al., 2025), which suggests the canonical HMM should also transfer well to OPM data. In the future, a combined canonical HMM incorporating data from Elekta, CTF and OPM devices is possible, which should improve the applicability of the canonical HMM to different modalities.

#### 4.5.2 EEG

Due to differences in the underlying signal being measured (magnetic field for MEG and electric field for EEG), MEG and EEG capture different aspects of brain activity and have different spatio-spectral sensitivities (Hari and Puce, 2023). Like MEG, EEG data can be source reconstructed to estimate activity within the brain. However, EEG data is typically noisier and has a lower channel density compared to MEG. Combined with the sensitivity to changes in the electromagnetic properties of the head, this often makes EEG data more difcult to source reconstruct (Hari and Puce, 2023). Leveraging information from parcellated MEG could help improve the inferences in EEG data. The canonical HMM approach proposed here aims to do this. In Figure 5, we see that adopting the canonical HMM allows us to identify MEG-like networks in the parcellated EEG data. This approach could facilitate studies of large-scale cortical networks using medium/high-density EEG. However, the user should note given the (current) canonical HMM was trained on Elekta MEG data that there can be systematic biases related to modality differences, which should be controlled for in subsequent analysis.

### 4.6 Fine Tuning

In scenarios where the dataset of interest is believed to be outside the distribution of the training data, i.e. the new boutique dataset contains features that the model was not trained on, there is an option to *fine tune* the canonical HMM. Examples include: data collected using a different scanner/modality; data that used an alternative preprocessing/source reconstruction; and data recorded during a task not in the training dataset. In fine tuning, we initialise the model parameters (initial state probabilities, transition probability matrix, and state covariances^3^) of a new HMM with the canonical HMM parameters. Then, the new HMM is trained for a short period on the boutique dataset. This training allows the new HMM to find the model parameters that best describe the boutique dataset. This allows us to benefit from the canonical HMM by transferring information from the larger dataset, which can help stabilise the inference and reduced computational demand, while identifying dataset-specific networks. However, in doing this we may lose the ability to compare networks across datasets.

The variational free energy (Bishop and Nasrabadi, 2006) can be used to determine whether the canonical HMM provides a good description of the boutique dataset. If the variational free energy for the boutique dataset is similar to the variational free energy of the training data, fine tuning is likely not required.^4^ The variational free energy for sessions in the training dataset are provided in the public code repository.

### 4.7 Number of States

An important choice when training an HMM is the number of states. Each specification for the number of states is a different model for the data. In this work, we use variational Bayesian inference to train the HMM, which involves optimising the *variational free energy* (Bishop and Nasrabadi, 2006). This objective function accounts for the number of states by adding a term to the training loss that penalises model complexity (i.e., a higher number of states). The variational free energy is an approximation for the *model evidence* (Friston et al., 2007), which can be used to compare different models and select one for subsequent analysis (Alonso and Vidaurre, 2023). The model with the lowest variational free energy is one that best fits the data whilst accounting for the number of states chosen. Figure S7A shows the variational free energy decreases with the number of states fitted to the Cam-CAN dataset (*N* = 621, rest and task). This indicates the more states we fit, the better we model the Cam-CAN data, and the optimum number of states according to this metric is above 40. Note, there are other clustering metrics, such as the silhouette score, which might be used to assess the quality of state assignments. These metrics are often based on distances between observed data points, which unfortunately do not reflect differences in the underlying covariance structure very well. Consequently, these clustering metrics do not identify clear optima in the number of states. This is illustrated in Figure S9 where we compare the silhouette score and variational free energy with simulated data.

How well we fit a particular dataset (as measured by the variational free energy) is not the only factor we consider when selecting the number of states. We wish to identify states that exist across a large population rather than modelling noise that is only present in one particular dataset. That is, we would like to accurately describe data from participants not seen when training the model. We seek states that can be *reproduced*^5^ in new data. A common approach for assessing this in neuroimaging studies is *split-half validation*^6^, where a model is trained on each half of the data independently and the similarity of the trained models is assessed. States that appear in both halves are reproducible. Figure S7 shows the split-half reproducibility of the states inferred on the Cam-CAN dataset.

Each HMM state corresponds to a spatio-spectral pattern of brain activity that arises when the state is active (power map). To assess the similarity between a model from each half of Cam-CAN, we calculate the average Pearson correlation between the power maps of each state (after re-ordering to find the best pairing). We observe a drop off in the average correlation between matched states from each half at around 20 states (Figure S7B). That is, we can identify 20 states that are common in two independent fits to separate halves of the Cam-CAN dataset (Figure S7C). Figure S8 shows the split-half reproducibility using a smaller subset of Cam-CAN (*N* = 50 and *N* = 16, rest and task). We see with less data, we cannot identify as many reproducible states.

By convention, it is common to select between 8 and 12 states (Vidaurre et al., 2018b; Cho et al., 2024; Gohil et al., 2024a; Higgins et al., 2021; Gohil et al., 2024b; van Es et al., 2023; Gohil et al., 2022; Quinn et al., 2018; Vidaurre et al., 2018a, 2016; Baker et al., 2014). We make parcel and sensor-level canonical HMMs with 4 to 16 states publicly available (see Section 6). The most common and recommended approach for selecting the number of states is to ensure any findings are robust to the number of states selected, i.e. the same effects such as group differences or network responses are present when a different number of states is chosen. This makes the exact choice for the number of states less critical.

### 4.8 Task-Relevance of the Canonical HMM Networks

In Figure 4, we studied the network activity in a working memory task. We can also see how the canonical HMM networks respond to the various tasks in the Cam-CAN dataset. The tasks included in Cam-CAN were a passive visual and auditory stimulus and a sensorimotor response (button press following a visuo-auditory stimulus). See (Shafto et al., 2014; Taylor et al., 2017) for further details regarding the tasks.

We can study the network-level response (Gohil et al., 2024b) to the various stimuli by epoching the state time courses around stimulus events. This is shown in Figure 8. We see multiple networks activate/deactivate in response to the stimuli. Four networks (states 1, 2, 9 and 10) show the same sequential response to the stimuli (Figure 8B). These networks correspond to higher-order cognitive function -this is discussed further in Section 4.9. We see networks that correspond to activity in unimodal primary cortices (Rolls, 2016): states 3 and 4 exhibit activity in the primary visual cortex; states 7 and 8 exhibit activity in the primary auditory cortex; and states 5 and 6 exhibit activity in the primary somatosensory/motor cortex. We can therefore associate these networks to the processing of primary sensory inputs.

**Figure 8:**
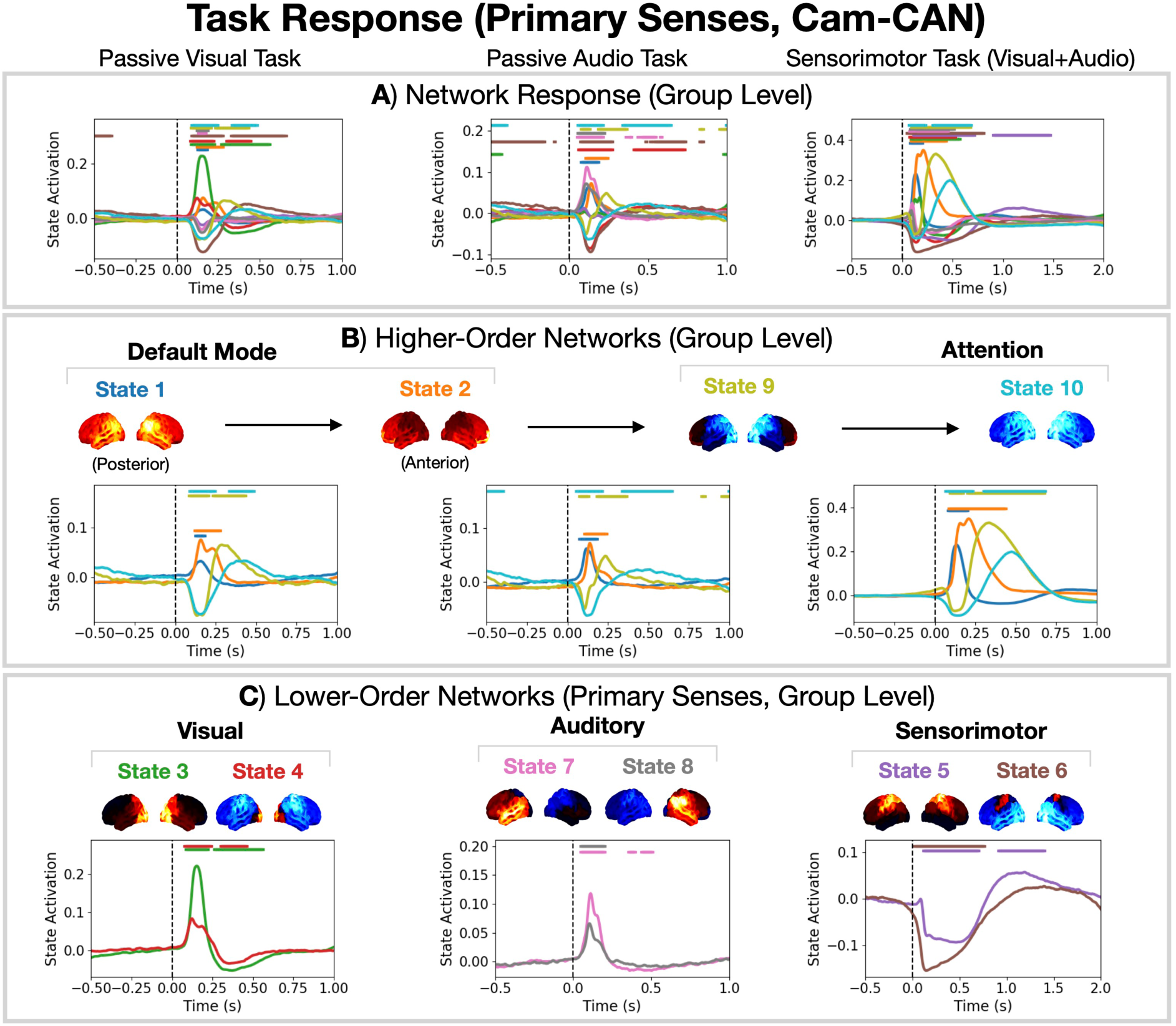
The canonical HMM networks provide a useful description for a variety of tasks. A) Network response (using the 10-state canonical HMM networks) to a passive visual task (left), passive audio task (middle) and visual/auditory sensorimotor task (right). See (Shafto et al., 2014; Taylor et al., 2017) for details regarding the tasks. B) Network response for the higher-order networks (states 1, 2, 9, 10). C) Network response for the lower-order networks corresponding to primary sense responses (states 3 and 4 for visual, states 7 and 8 for auditory, and states 5 and 6 for sensorimotor). Horizontal bars indicate time points with *p*-value < 0.05, which were calculated using sign-flip GLM permutations taking the maximum statistic (COPE) across time and states to control for multiple comparisons.

The canonical HMM provides a flexible basis set that can describe a variety of cognitive functions in both task and rest data. Adopting the canonical HMM for task analysis can be useful in the study of disease. Particularly if the symptoms of disease only arise in task (Casagrande et al., 2023).

The natural question that arises is whether the HMM networks themselves are modified by task. Previous work suggests this may be the case. For example, Colenbier and colleagues showed task-induced changes to (static) functional networks in MEG occur and that these changes improve our ability to identify individuals (Colenbier et al., 2023). However, depending on the size of the task-induced changes, the canonical HMM may or may not be a good approximation for functional networks in task. In the MEGUK *N*-back task, we see very similar networks are inferred when training a model from scratch and applying the canonical HMM (Figure 4A and B respectively). This suggests the task-induced changes are small (in *N*-back task at least) and the canonical HMM is a good approximation. Note, the calculation of the multitaper spectra in the post-hoc analysis is based on the boutique data. Therefore, we are still able to identify and study task-induced changes in the networks even if we use the canonical HMM. Furthermore, if we wish to adapt the HMM to the boutique task dataset, then we have the option to fine tune the canonical HMM. Understanding how task affects the canonical HMM networks and whether fine tuning is necessary is an interesting future study.

### 4.9 Correspondence Between MEG and fMRI Networks

Here, we compare the spatial power maps of the canonical HMM states to the fMRI networks presented in (Yeo et al., 2011), which are shown in Figure 9A. We divide the fMRI networks into two types: those that correspond to *lower-order* cognitive function (primary sensory input, networks 1 and 2) and those that correspond to *higher-order* cognition function, i.e. functions that go beyond primary sensory input (networks 3-7) (Van Den Heuvel and Pol, 2010; Smitha et al., 2017). Higher-order cognitive functions include processes such as executive control, memory and attention (Smitha et al., 2017; Kiely, 2014). Correlating the HMM MEG state power maps with the fMRI networks, we see the MEG networks can also be divided into those that correspond to higher-order and lower-order cognitive function (Figure 9).

**Figure 9:**
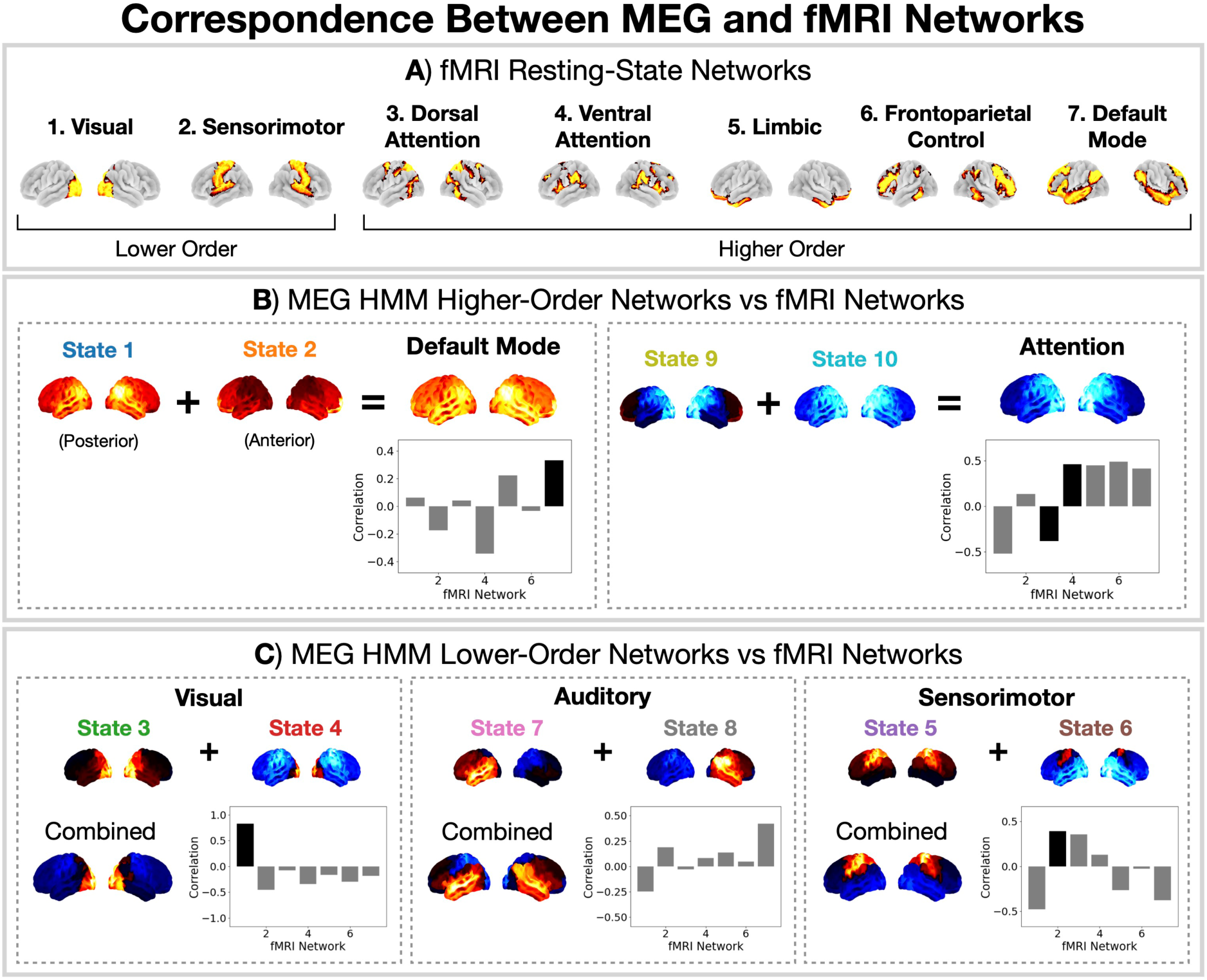
Correspondence between the canonical MEG networks and fMRI resting-state networks. A) Seven fMRI resting-state networks identified by (Yeo et al., 2011). FMRI networks 1 and 2 correspond to primary sensory input (*lower-order* cognitive function) and networks 3-7 correspond to *higher-order* cognitive function. B) Correlation between the higher-order MEG networks (default mode, left; attention, right) and the fMRI networks. C) Correlation between the lower-order networks (primary sensory input) and fMRI networks. In black we highlight the pairing of the MEG and fMRI networks.

Of particular interest is the *default mode* network from fMRI (network 7 in Figure 9A), which is associated with internally oriented cognitive functions (Raichle, 2015) and tends to activate in the resting state (Van Den Heuvel and Pol, 2010). In MEG, it was found that two HMM states correspond to a default mode network: a posterior and an anterior network (canonical HMM states 1 and 2 respectively) (Vidaurre et al., 2018b). This division is due to the higher temporal resolution provided by MEG (milliseconds) compared to fMRI (seconds). Combining these two HMM states we find a spatial pattern of power that correlates with the fMRI default mode network (Figure 9B).

A well-known finding in the fMRI literature is the anti-correlation between the dynamics of the default mode network (7 in Figure 9A) and the dorsal attention network (3 in Figure 9A) (Fox et al., 2005). The corresponding networks in MEG that have anti-correlated dynamics to the default mode network are states 9 and 10. This was previously shown in (Baker et al., 2014) and is reflected by the lack of transitions between these states (Figure 2D). Correlating the combination of states 9 and 10 to the fMRI networks we see strong associations with all of the higher-order fMRI networks. Therefore, based on the spatial pattern of activity alone these states cannot be associated to a single fMRI network. However, based on the anti-correlation in terms of dynamics and the suppressed posterior-*α* (8-12 Hz) activity exhibited by the states, which is common marker for spatial attention (Gould et al., 2011), we consider states 9 and 10 to be attention networks in MEG.

Turning to the lower-order fMRI networks, we can see MEG states 3 and 4 correspond to the visual network (Figure 9C) and MEG states 5 and 6 correspond to the sensorimotor network (Figure 9C). In MEG, we are also able to identify a network (states 7 and 8) that correspond to auditory inputs, which is not present in the fMRI networks from (Yeo et al., 2011).

## 5 Conclusions

We present a canonical HMM framework that captures reproducible, fast-switching electrophysiological networks from a large cohort of healthy individuals (Cam-CAN; 1849 sessions, *N*=621). The released models (4-16 states, parcel and sensor-level) provide a common reference that drastically reduces computational cost of performing an HMM analysis on M/EEG. This also improves the stability and comparability of network inferences across boutique MEG/EEG studies. We demonstrate the approach on multiple example datasets (rest and task), showing good agreement with conventional, dataset-specific HMMs. Important practical constraints include: matching the preprocessing, source reconstruction and parcellation choices (sampling rate, orthogonalisation, sign-flipping, PCA/TDE parameters). Alternatively, a sensor-level model can be used provided the data was collected with the same scanner type. Future work will extend the canonical HMM to include additional scanners/modalities, a more diverse set of tasks including higher-order cognitive processes, and explore fine-tuning strategies. By providing these models and example code as an open resource, we aim to make HMM analyses in M/EEG more accessible, reproducible and comparable across studies.

## 6 Data and Code Availability

All the datasets used in this work are publicly available (see Section 2.1). Code and examples scripts for processing M/EEG data and applying the canonical HMM have been made publicly available here: https://github.com/OHBA-analysis/Canonical-HMM-Networks. As larger and more diverse datasets become available this resource will be actively updated. This includes providing canonical HMMs with differing model orders, different parcellations, different sensor layouts and training on larger datasets including other modalities, such as OPM and EEG data.

## 7 Author Contributions

CG: Conceptualisation; Methodology; Software; Validation; Formal Analysis; Investigation; Data Curation; Writing -Original Draft; Writing - Review & Editing; Visualisation. RH: Software. CH: Visualisation. MWJE: Validation. AJQ: Conceptualisation. DV: Writing - Review & Editing. MWW: Conceptualisation; Methodology; Writing - Review & Editing; Supervision.

## Footnotes

1 For the Elekta MEG datasets, we applied these steps to the publicly available MaxFiltered data. For the other datasets, we used the publicly available raw sensor recordings.

2 Note, the Markovian constraint does not prevent us from inferring long-range temporal dependencies in real data. Although the HMM generative model only considers the previous time point, the real data may have been generated from a process with a longer memory. This information is contained in the observed data. In calculating the state probabilities (posterior) we can infer long-range dynamics in the states from the data directly despite the short memory of the HMM.

3 We assume the state means are zero.

4 This is an approach commonly used in machine learning where the loss is calculated on a held out test dataset and compared to the training loss.

5 Here, we refer to the same concept as *generalisation*, which is the term used in the machine learning literature.

6 Equivalent to 2-fold cross validation.

