## Supplementary Information for "Canonical Hidden Markov Model Networks for Studying M/EEG"

### Supplementary Information (SI)

Table S1: Hyperparameters used to train the Hidden Markov Model in `osl-dynamics`.

| Hyperparameter | Value |
|---|---|
| `batch_size` | 32 |
| `sequence_length` | 400 |
| `learning_rate` | 0.001 |
| `n_epochs` | 10 |
| `optimizer` | `adam` |
| `observation_update_decay` | 0.1 |
| `trans_prob_update_delay` | 5 |
| `trans_prob_update_forget` | 0.7 |

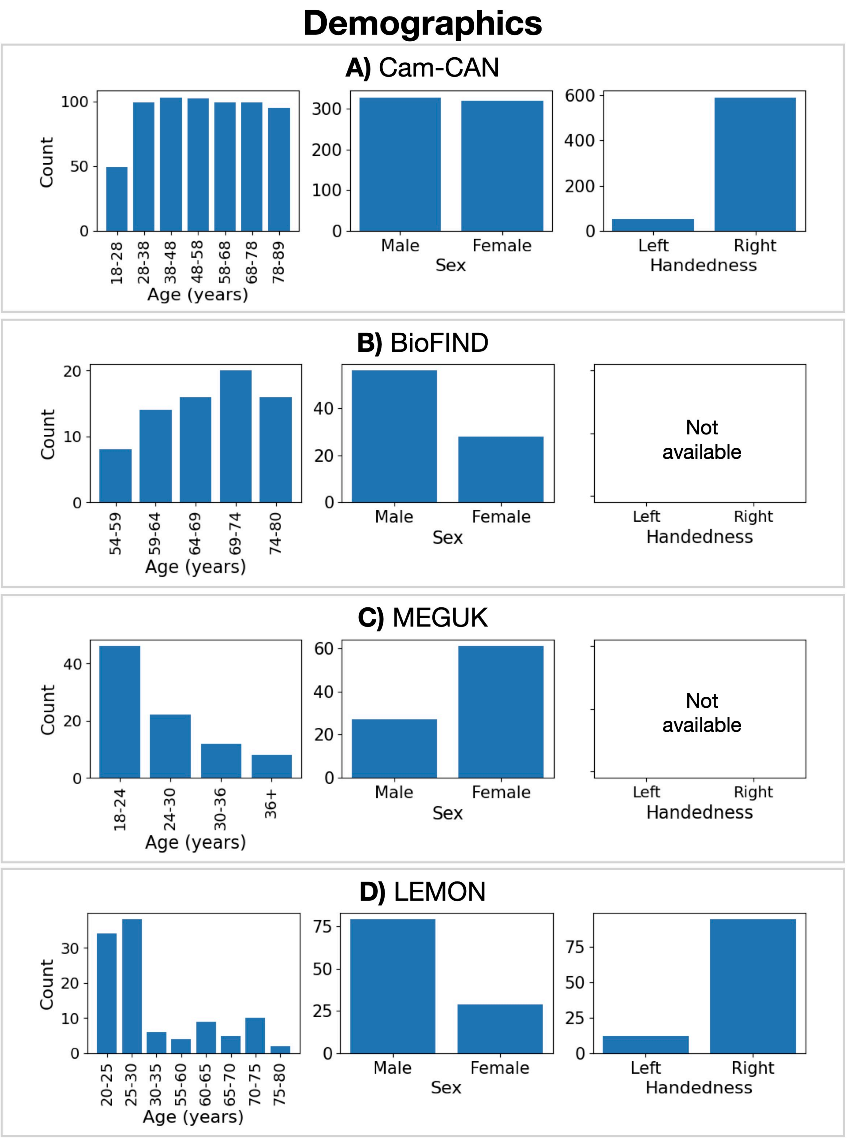


Figure S1: **Demographics for the datasets studied in this work**. Distribution of ages (left); sex (middle) and handedness (right).

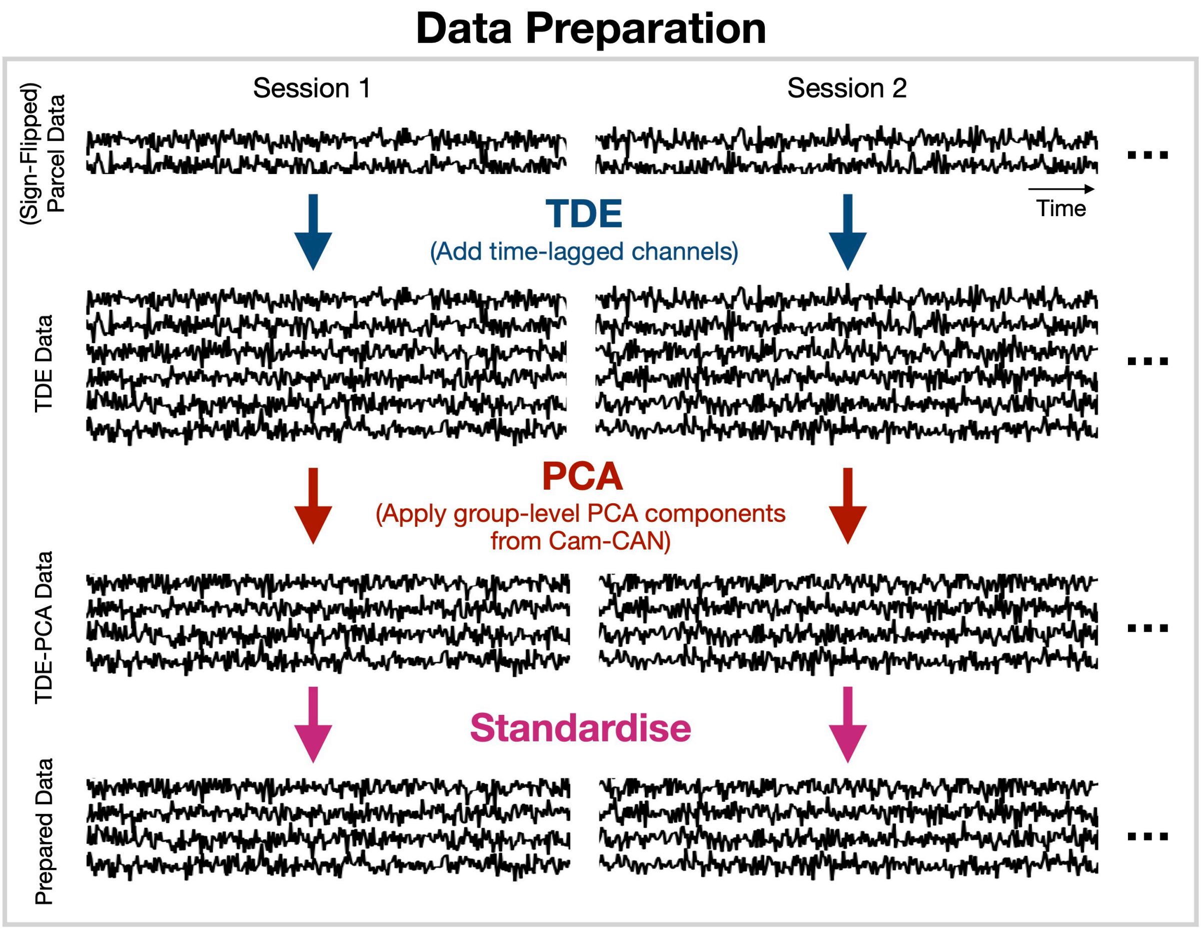


Figure S2: **Data preparation**. For each session, we first time-delay embed (TDE) the parcel time courses (sign-flipped to match the Cam-CAN template session). Following this, we apply the group-level Cam-CAN principal component analysis (PCA) components (calculated on the TDE Cam-CAN parcel data) to reduce the dimensionality of the TDE data.

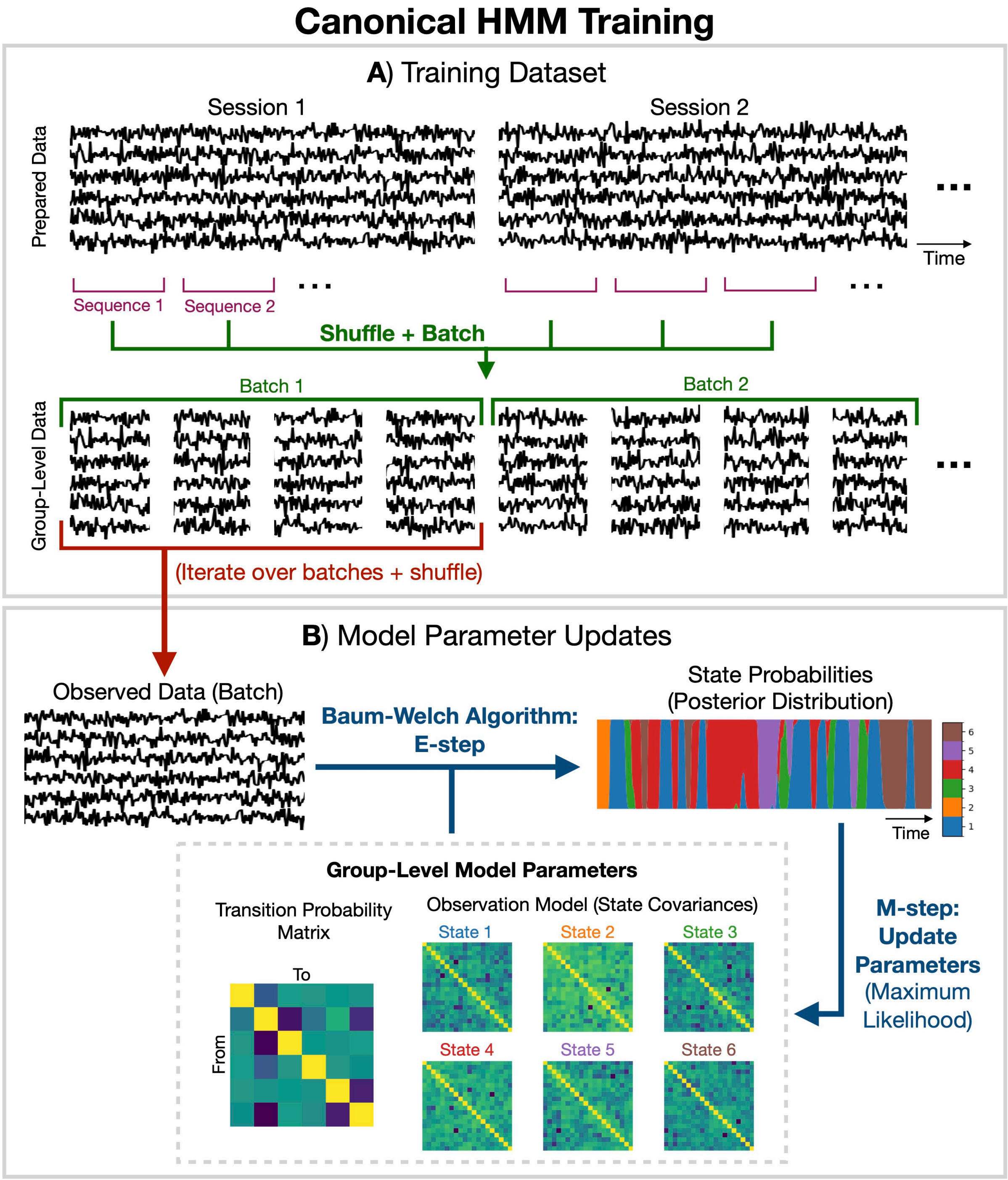


Figure S3: **Canonical HMM training**. A) Handling of the training data. First we split the time series into sequences, then shuffle, batch, and shuffle again. B) Iterative EM algorithm to inference the state probabilities and update the group-level model parameters.

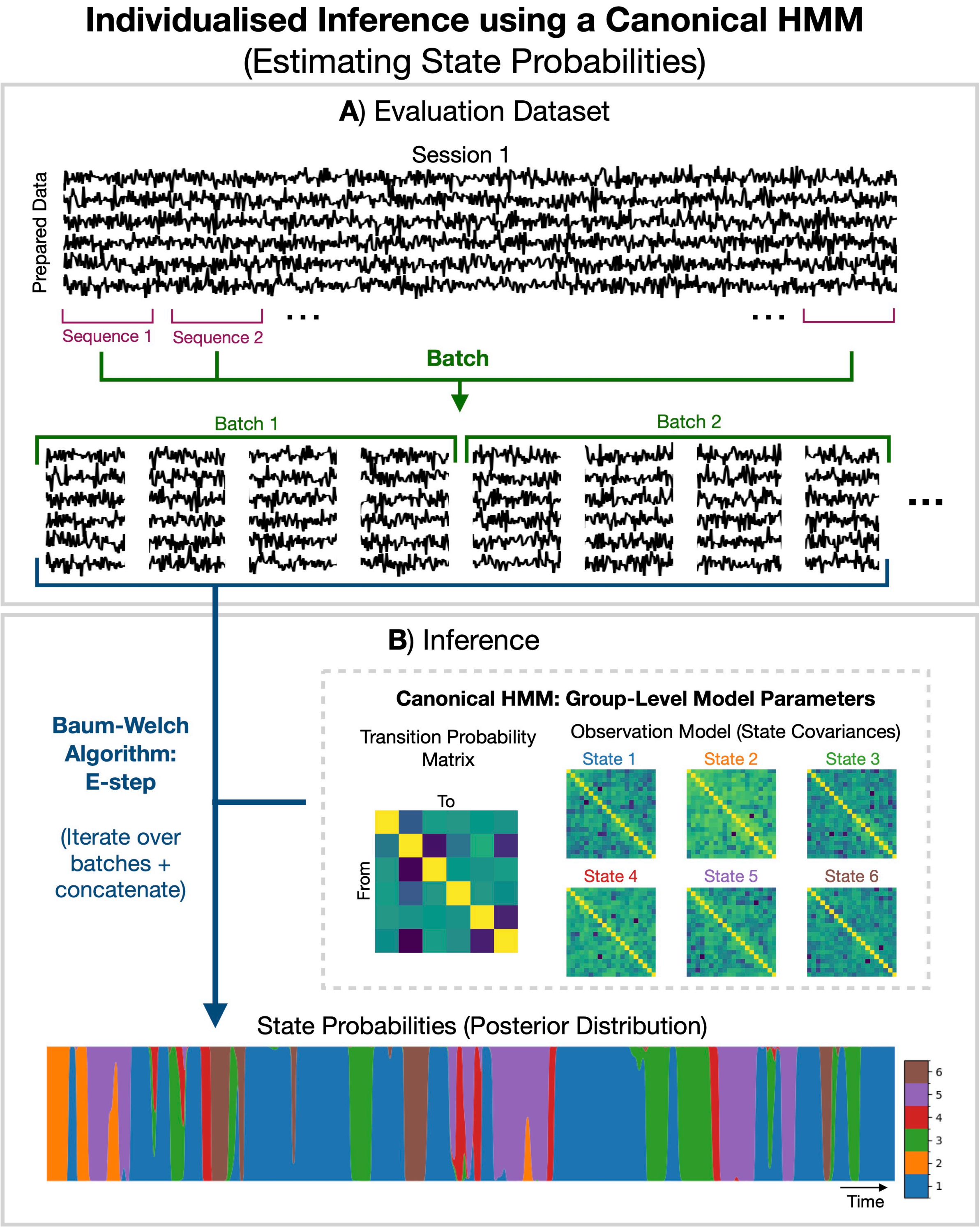


Figure S4: **Individualised inference using a canonical HMM**. A) Handling of the evaluation data. We separate the time series in sequences and batch. B) Inference of state probabilities for the evaluation data based on the canonical HMM group-level parameters.

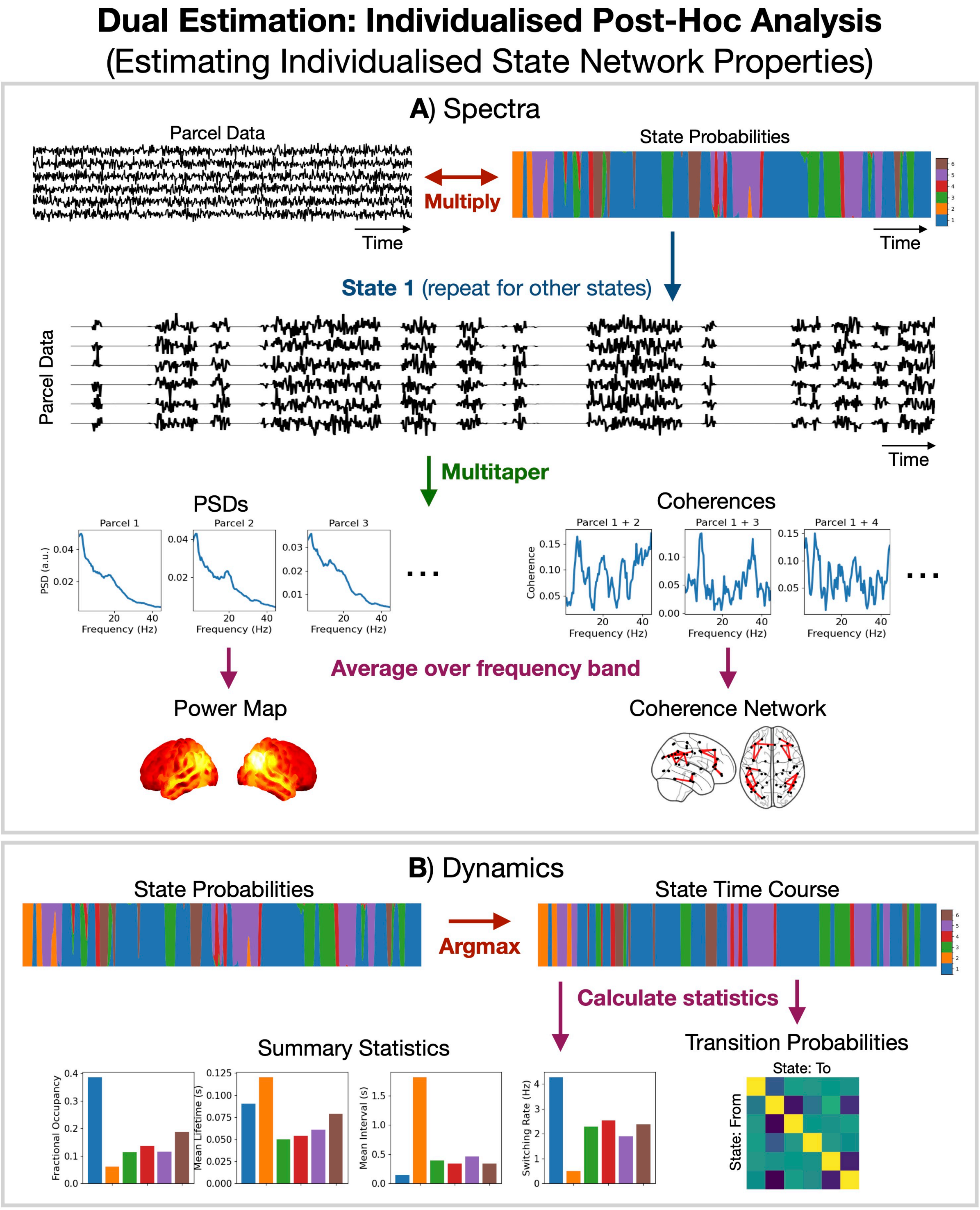


Figure S5: **Individualised post-hoc analysis (dual estimation)**. A) Calculation of session and state-specific multitaper spectra, power maps and coherence networks. B) Calculation of summary statistics for dynamics.

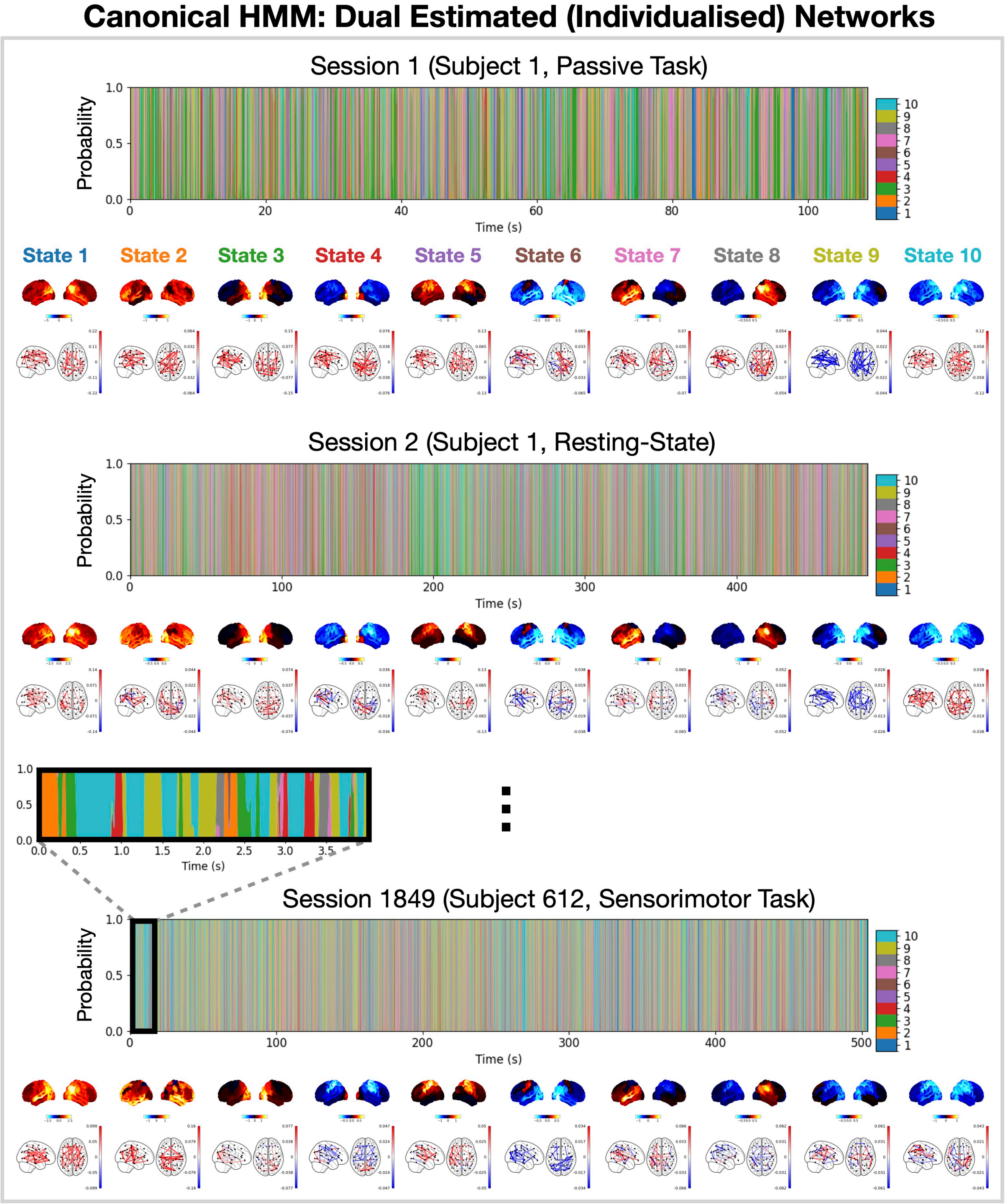


Figure S6: **Individualised networks from the 10 state canonical HMM**. State probability inference was performed on each session from the Cam-CAN dataset.

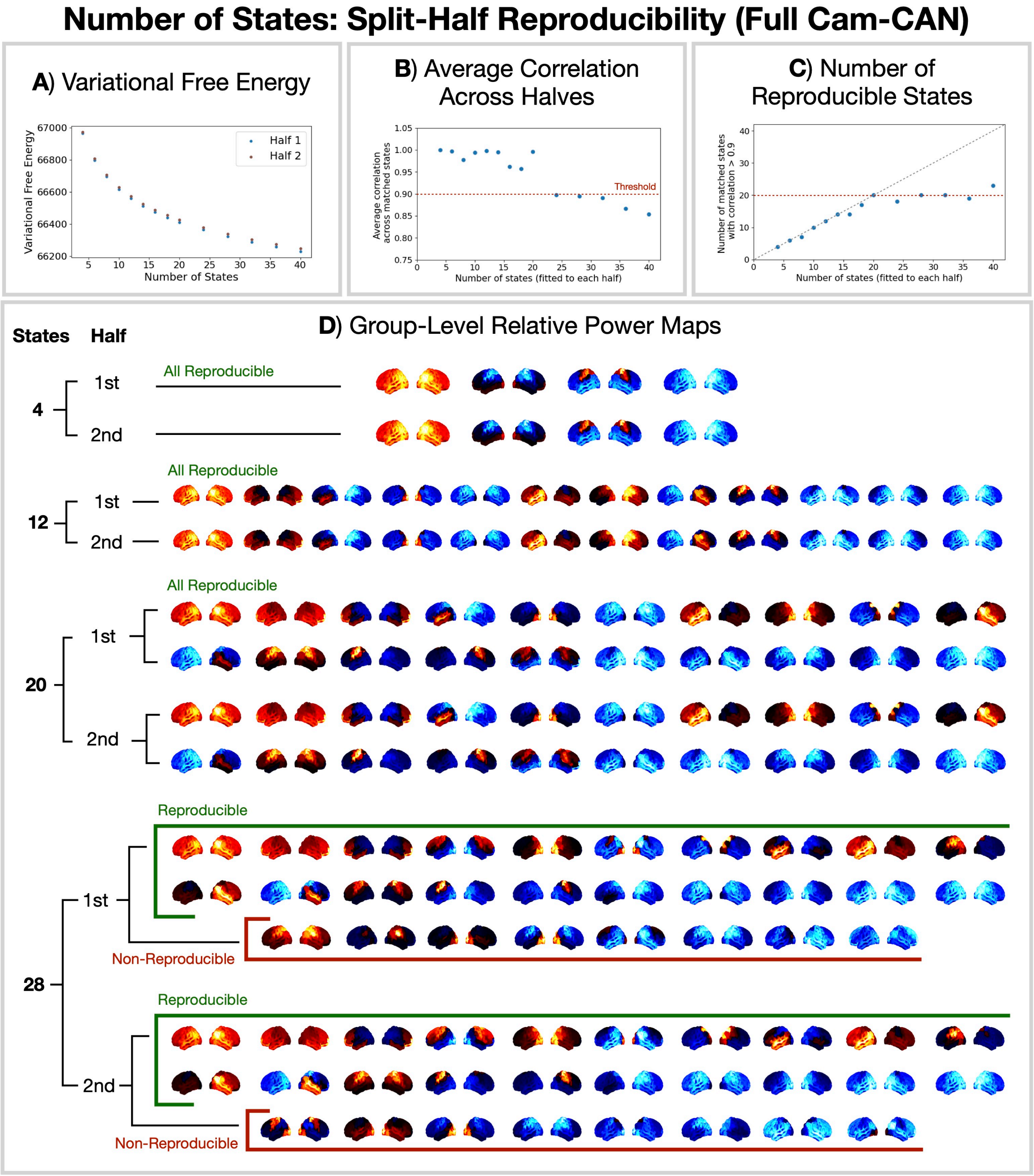


Figure S7: **Up to 20 HMM states can be reproducibly inferred with the Cam-CAN MEG dataset**. A) Final variational free energy averaged over 10 runs on each half vs number of states. B) Average Pearson correlation between power maps from each half vs number of states. C) Number of states from each half that had a correlation above 0.9 vs number of states. When comparing the halves, the states were re-ordered to maximise pairwise correlation between the power maps using the Hungarian algorithm. D) State power maps (1-45 Hz) relative to the mean across states for differing numbers of HMM states for each half.

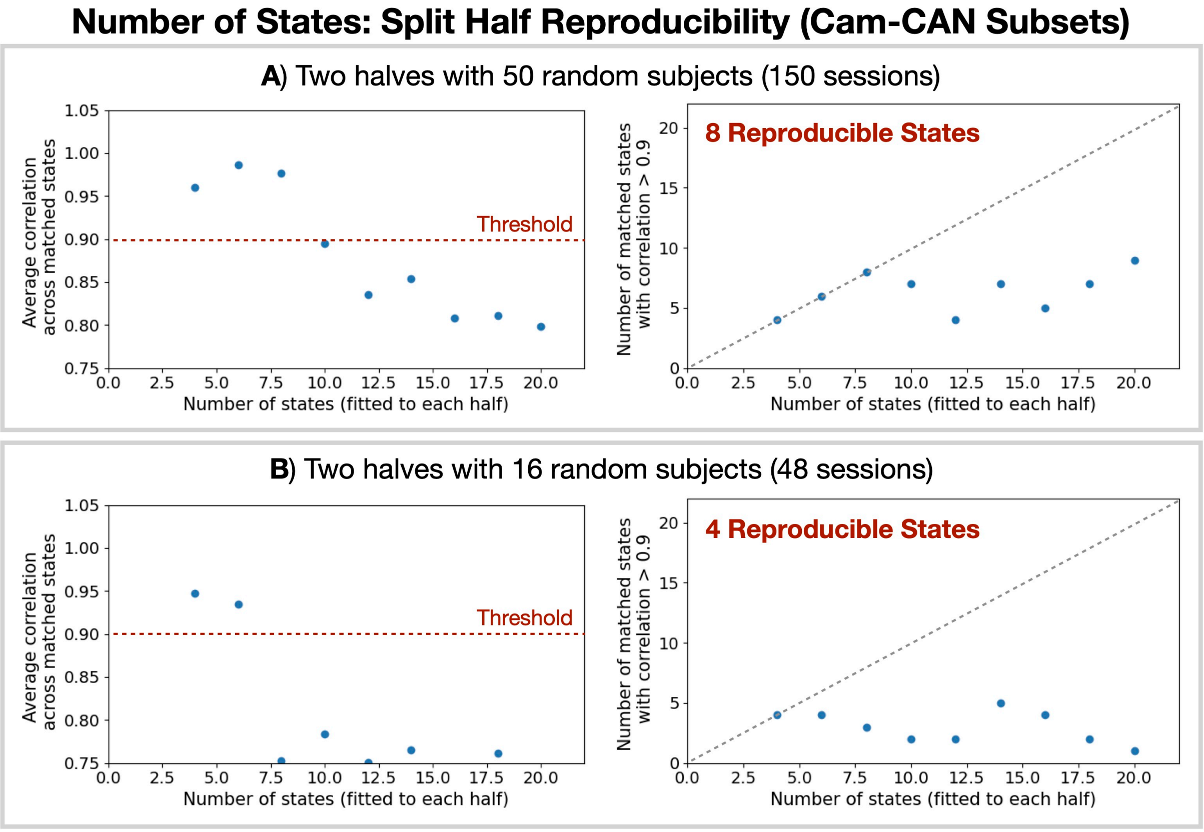


Figure S8: **Split half reproducibility with subsets of Cam-CAN**.

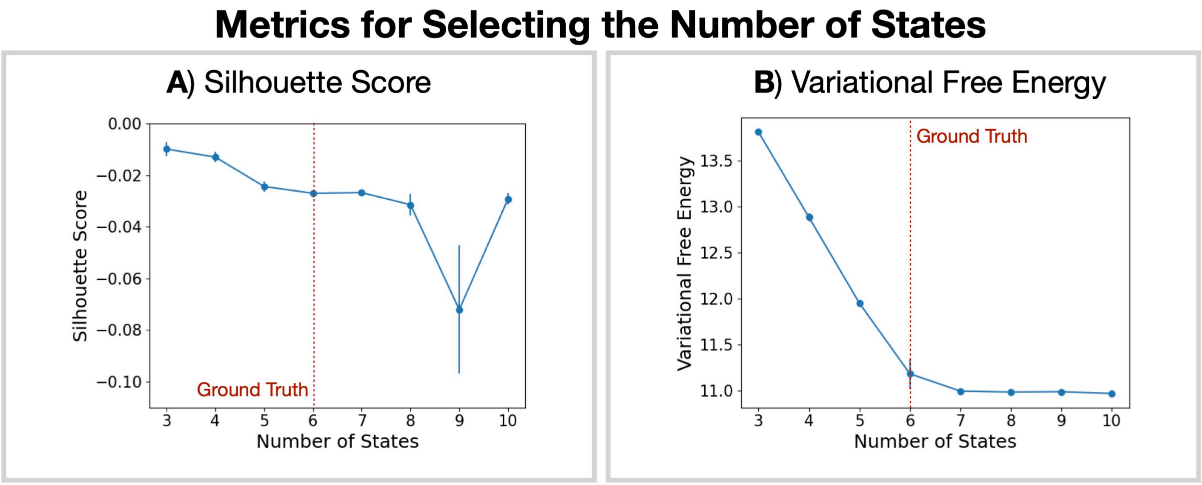


Figure S9: **Metrics for selecting the number of states**. A 6 state HMM was simulated with zero and random covariances. A transition probability with stay probability 0.9 and equal probability of transitions to any other state was used. Initial state probabilities were a uniform distribution. We trained an HMM with 3-10 states (5 runs) on the simulated data and calculated the silhouette score (A; higher is better) and variational free energy (B; lower is better). Error bars show the standard error on the mean. We see the variational free energy plateaus at the correct number of states, however, the silhouette score does not identify a clear optimum. The negative values of the silhouette score suggests the data points are misassigned.
